# Patterns, Profiles, and Parsimony: dissecting transcriptional signatures from minimal single-cell RNA-seq output with SALSA

**DOI:** 10.1101/551762

**Authors:** Oswaldo A. Lozoya, Kathryn S. McClelland, Brian Papas, Jian-Liang Li, Humphrey H-C Yao

**Affiliations:** Genomic Integrity & Structural Biology Laboratory, National Institute of Environmental Health Sciences, Research Triangle Park, North Carolina, USA; Reproductive and Developmental Biology Laboratory, National Institute of Environmental Health Sciences, Research Triangle Park, North Carolina, USA; Integrative Bioinformatics Support Group, National Institute of Environmental Health Sciences, Research Triangle Park, North Carolina, USA; KSM now at: Section of Developmental Genomics, Laboratory of Cellular and Developmental Biology, National Institute of Diabetes and Kidney and Digestive Diseases, National Institutes of Health, Bethesda, Maryland, USA

## Abstract

Single-cell RNA sequencing (scRNA-seq) technologies have precipitated the development of bioinformatic tools to reconstruct cell lineage specification and differentiation processes with single-cell precision. However, start-up costs and data volumes currently required for statistically reproducible insight remain prohibitively expensive, preventing scRNA-seq technologies from becoming mainstream. Here, we introduce single-cell amalgamation by latent semantic analysis (SALSA), a versatile workflow to address those issues from a data science perspective. SALSA is an integrative and systematic methodology that introduces matrix focusing, a parametric frequentist approach to identify fractions of statistically significant and robust data within single-cell expression matrices. SALSA then transforms the focused matrix into an imputable mix of data-positive and data-missing information, projects it into a latent variable space using generalized linear modelling, and extracts patterns of enrichment. Last, SALSA leverages multivariate analyses, adjusted for rates of library-wise transcript detection and cluster-wise gene representation across latent patterns, to assign individual cells under distinct transcriptional profiles via unsupervised hierarchical clustering. In SALSA, cell type assignment relies exclusively on genes expressed both robustly, relative to sequencing noise, and differentially, among latent patterns, which represent best-candidates for confirmatory validation assays. To benchmark how SALSA performs in experimental settings, we used the publicly available 10X Genomics PBMC 3K dataset, a pre-curated silver standard comprising 2,700 single-cell barcodes from human frozen peripheral blood with transcripts aligned to 16,634 genes. SALSA identified at least 7 distinct transcriptional profiles in PBMC 3K based on <500 differentially expressed Profiler genes determined agnostically, which matched expected frequencies of dominant cell types in peripheral blood. We confirmed that each transcriptional profile inferred by SALSA matched known expression signatures of blood cell types based on surveys of 15 landmark genes and other supplemental markers. SALSA was able to resolve transcriptional profiles from only ∼9% of the total count data accrued, spread across <0.5% of the PBMC 3K expression matrix real estate (16,634 genes × 2,700 cells). In conclusion, SALSA amalgamates scRNA-seq data in favor of reproducible findings. Furthermore, by extracting statistical insight at lower experimental costs and computational workloads than previously reported, SALSA represents an alternative bioinformatics strategy to make single-cell technologies affordable and widespread.

## Introduction

Next-generation sequencing technologies are transforming how biologists characterize the molecular features of organogenesis and the composition of heterogeneous tissues; among them, RNAseq is one of the most widely adopted modalities [1-3]. RNAseq can be used to understand how intricate transcriptional networks regulate cell fate determination and lineage specification during organogenesis, development, and disease [4-6] – by profiling cell lines, sorted primary cells, and bulk tissues [7-9]. Yet, although bulk RNAseq experiments have sufficed to determine the benchmark of gene expression signatures that underlie whole-organ physiology, they have always been inadequate to distinguish cell-type specific transcriptional dynamics. Understanding the inherent variety present in mixed pools of individual constituent cells, all different from each other at any given time, based on their location within tissues, or stage during lineage specification, remains an outstanding question.

Resolving spatiotemporal relationships in the transcriptional profile of tissues during the most dynamic periods, such as during embryonic development, presents a major challenge. Splitting heterogeneous cell mixtures (from tissues) into subsets and performing bulk RNAseq in each subpopulation of collected cells separately has provided rich new details, yet targeted profiling of sorted cell populations remains insufficient [10]. Importantly, extracting transcriptional signatures of targeted cell populations is only possible if relevant markers are known in advance. Even then, it is well appreciated that artificially pooling cells by sorting risks relying on the faithful and stable expression of lineage markers, which may be inconsistent with their physiological reality, and may distort the transcriptional dynamics of cell populations *in vivo* [11].

Single-cell transcriptomics circumvents many of these obstacles, and recent breakthroughs in RNAseq technologies now allow reconstruction of cell lineage specification processes in embryonic tissues at the level of individual cells [12-18]. A diverse catalogue of single cell RNAseq (scRNA-seq) platforms and workflows are currently available to interrogate transcriptomes from individual cells. Using bioinformatic tools, individual cells are sorted and classified by their gene expression similarities, and the identity of each cell is deduced based on their transcriptional and functional ontology [19-22]. Broadening the scope of samples that can undergo scRNA-seq workflows, multiple groups have introduced chemical treatments for freshly excised specimens, akin to whole-tissue mounting, that allow long-term storage of specimens whole and conserving their endogenous transcripts. These modifications make a wider range of collected and banked tissues, including clinical samples, compatible with scRNA-seq protocols designed for freshly excised and dissociated specimens. Some of these treatments include methanol fixation [23], actinomycin D quenching of RNA polymerases activity [24], and free-amine cross-linking with dithio-bis(succinimidyl propionate) (DSP) [25].

With the numerous customizable single-cell techniques and off-the-shelf modalities becoming available come new challenges to researchers around the analysis of scRNA-seq data. To date, there is neither consensus, nor uniformity, about how best to evaluate experimental designs that compare independent biological specimens, or, how to integrate data across biological replicates. Different commercial platforms offer a range of pre-packaged analytical pipelines, workflows and statistical models to handle scRNA-seq data, but they typically can only analyse combined outputs of multiple sequencing runs as a single all-at-once dataset. This approach is suboptimal because it disregards any statistical or batch mismatches inherent to experimental replication. Moreover, data obtained from competing platforms currently does not translate seamlessly between their respective post-processing workflows, making it challenging to integrate and compare data generated by different scRNA-seq platforms. Some elegant solutions to integrate data from separate specimens have become available recently [26,27], however all these approaches require that datasets must be derived from technologies that all accomplish single-cell barcoding using one of three broad strategies to physically sift individual cells: plucking [16,28-35], droplet-based encapsulation [14,15,36,37], or split-pooling [13,18]. Little information exists that explicitly tackles how to reconcile scRNA-seq data from replicate specimens where libraries were synthesized with technologies based on different cell sifting strategies.

In this work, we introduce a versatile workflow, named single-cell amalgamation by latent semantic analysis (SALSA), that unifies parametric criteria for barcode (and transcript) filtering from raw scRNA-seq data. To evaluate SALSA we benchmarked its cell type discriminative power using a publicly available and widely regarded “silver” standard, the single-run 10X Genomics frozen 3K PBMC data obtained with the commercially available Chromium platform [36]. In all, we show that SALSA successfully extracts replicable transcriptional signatures, and reproduces functional ontologies by offering a new approach to scRNA-seq data mining. By extracting latent variable signatures from ultra-sparse scRNA-seq datasets, SALSA offsets the cost of performing biological replication through savings in sequencing output. These advances set a path forward to allow multi-platform confirmatory testing, orthogonal hardware comparisons, and overall technological dissemination to be carried out with limiting resources.

## Results

### Frequentist expression matrix reduction via mixture model-guided focusing

A critical phase in scRNA-seq data analysis preceding statistical inferential testing between genes or cell barcodes, is curation of gene-cell (or cell-gene) matrices. When performed systematically, curation of gene-cell matrices can lay down a path towards statistical robustness and reproducibility, making comparisons of scRNA-seq datasets across multiple samples more reliable. This essential first step aims to reproducibly infer which detected barcodes correspond to single cells, and which unique molecular identifiers (UMIs) align to variable genes in a reference genome, in datasets generated using droplet-based encapsulation or split-pooling techniques alike (Fig.1A). For SALSA, we performed gene-cell matrix curation through a parametric frequentist approach, which shifts focus away from net UMI coverage and towards the uneven apportionment of UMIs between artefactual, single-cell, and multi-cell barcodes (Fig. 1B).

**Figure 1.**
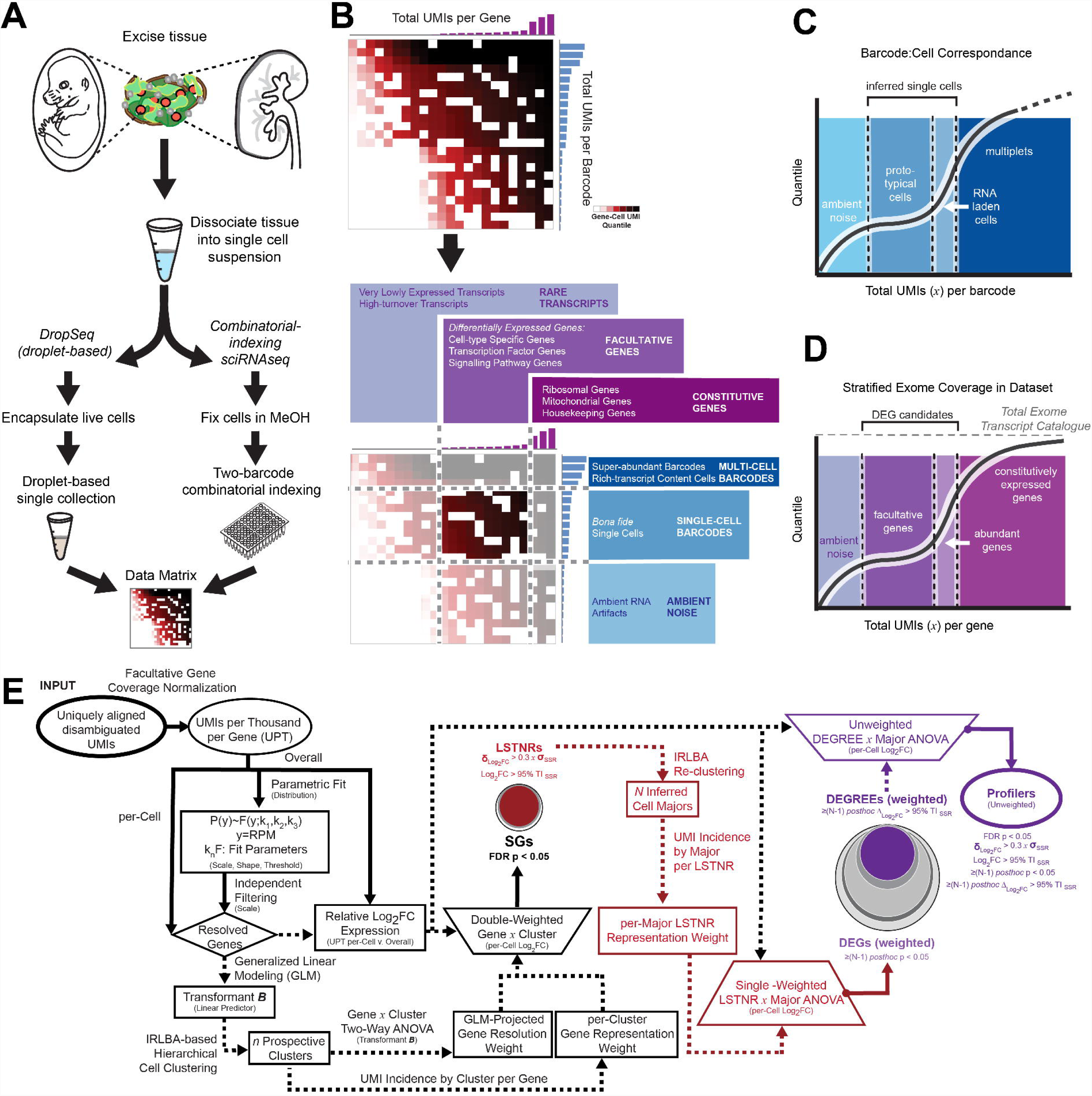
Implementation of the SALSA workflow. (A) To assemble a gene-cell expression matrix for thousands of cells in a single run by scRNA-seq, biological specimens are dissociated into single cell suspensions, partitioned for barcoding and adapterization by droplet encapsulation (e.g. DropSeq) or split-pooling technologies (e.g. sci-RNAseq), and sequenced with short-read high-throughput SBS instrumentation. (B) Matrix focusing for scRNA-seq expression matrices. Sorted count data from scRNA-seq experiments exhibits transitions in total UMI counts per barcode, reminiscent of distinct regimes of UMI density between background (ambient noise), single-cell, and multi-cell barcodes; total UMI counts per gene exhibit an analogous profile, with distinct regimes between rare, facultative, and constitutively expressed genes. Latent patterns of expression within gene-cell matrices are most discriminative at the intersection of facultative genes and single-cell barcodes regimes, referred to as the focused expression matrix. To infer coverage regimes per barcode (C) and per gene aligned (D) from the raw gene-cell expression matrix, total UMI count data are fit to a 2-component mixture probabilistic parametric model; regime thresholds are defined systematically from estimated scale and shape parameters. (D) Stratified differential expression analysis starting from a focused expression matrix in SALSA. The flow chart depicts transformations used in SALSA towards generalized linear modelling (GLM) of expression data, and statistical criteria to demarcate gene strata with increasing levels of prospective experimental reproducibility among facultative genes: statistically significant genes (SGs), leveraged signal-to-noise-ratio genes (LSTNRs), differentially expressed genes (DEGs), DEGs with reproducible expectation estimates (DEGREEs), and Profiler genes.

We reasoned that ascribing single-cell status to a barcode is driven by context: single-cell barcodes are more like each other than they are to ambient RNA or multi-cell barcodes. As the number of UMIs per barcode is commensurate with the total mRNA molecules each barcode was appended to during cDNA synthesis, a “single-cell regime” based on UMI totals should exist and be recognizable from artefacts (a form of rank similarity) and conserved at different sequencing depths (a form of scale invariance). Together, these assumptions of rank similarity [38,39] and scale invariance [40] are amenable to parametric fitting, particularly in the context of extreme value probabilistic models. Such models have found broad relevance in diverse research fields including the informational sciences of computational optimization [41], and financial forecasting in econometrics [42].

In the context of scRNA-seq, there are at least two “extreme” scenarios to consider in exploiting the features of extreme value theory: multi-cell barcodes are “rare events” relative to single-cell barcodes on the high-end of UMI coverage; and single-cell barcodes are “rare” relative to artefacts at low UMI coverages (Fig. 1C). We approached these two conditions simultaneously using a two-component model which we call the P_C_-P_D_ mixture model. Our model consists of a composite distribution that couples cases in which: a) barcoded artefacts and single-cells are both detected in similar numbers and differ by their total UMI counts, but share a common “floor” of contaminant UMI counts that is spread uniformly across the library (Frechét-like distribution, akin to <u>c</u>ombinatorial based scRNA-seq techniques where “noise lifts barcodes”); and b) barcoded artefacts, carrying low total UMI counts, are substantially more numerous than single-cell barcodes (Weibull-like distribution, akin to droplet-based scRNA-seq techniques where “noise gets barcodes”). Further details on the statistical treatment and parameterization of count-level data with the P_C_-P_D_ mixture model are available in the Supplementary Methods section.

To be implemented, the P_C_-P_D_ mixture model requires that gene-cell matrices be aggregated to compute empirical cumulative distribution functions (eCDFs) for total UMIs per detected barcode and total UMIs per aligned gene. This is followed by quantile regression of best-fit parametric scale and shape factors, and a partial contribution rate from one of the two constituent P_C_ and P_D_ distributions. Parametric scale factors of the P_C_-P_D_ mixture model are used to estimate critical inflection points at transition regions in the eCDF between low, transient, or maximal UMI counts per barcode, whereas parametric shape factors gauge the steepness of these transitions. Thus, when used in combination, P_C_-P_D_ parametric factors help demarcate the range of UMI coverages corresponding to single cells (Fig. S1). Using a similar logic, the same frequentist approach can be used to distinguish facultative genes from rare or constitutively expressed ones (Fig. 1C and D). From here on in the workflow, only data from “best-guess” single-cell barcodes, trimmed to retain facultative genes exclusively, is needed for downstream differential expression analysis of scRNA-seq datasets (Fig. 1E).

### The SALSA workflow

At its core, the SALSA methodology examines total UMI counts to infer transitions in UMI coverage between artefactual (baseline), single-cell (signal) and multi-cell (overshoot) barcodes using a parametric frequentist approach (Fig. 1C and D). SALSA then prioritizes information from *facultative* genes (those most likely to vary between individual barcodes) and projects the underlying single-cell × facultative gene UMI subset onto an imputable eigenvalue problem of linearized and normalized transformations of expression scores. To achieve this, SALSA calculates “bulk” expression levels of each facultative gene (i.e. all single cells added together), then uses them as a bulk-wise “reference mean” for generalized linear modelling (GLM) of single-cell expression levels [43]. With a normalizing and linearizing transformation at hand, SALSA exploits GLM transformants of single-cell expression levels to infer an *a priori* grouping of single-cells and facultative genes via a singular value decomposition (SVD)-driven implicitly restarted Lanczos bidiagonalization algorithm (IRLBA) coupled with Euclidean hierarchical clustering (Ward’s method) [44]. Next, to perform differential expression analysis, SALSA estimates log-fold expression changes in facultative genes per cell against the overall “reference mean” from the collective of single-cell barcodes. This is followed by examination of the overall degree of dispersion in log-fold expression from single cells around the mean of their prospective clusters using signal-to-noise ratio (SNR) benchmarking to 95% tolerance interval (95% TI_SSR_) of log-fold expression residuals. Significant genes (SGs) are inferred between prospective clusters using double-weighted multivariate analyses for both resolution of mean gene coverage (such as in the LSTNR method [45]), and gene representation rates within each of the prospective clusters. Then, SALSA identifies leveraged signal-to-noise ratio genes, or LSTNRs; these genes are the subset of SGs whose mean log-fold expression SNR>1 in at least one prospective cluster. Lastly, SALSA refines single-cell cluster assignment into inferred cell majors by performing *a posteriori* IRLBA clustering of GLM-transformed expression scores based only on the curated LSTNR genes. An abbreviated schematic of the SALSA workflow is outlined in Fig. 1E.

Experimentally, capturing a transcript that encodes a *bona fide* protein marker known as a hallmark representative of a population from a single cell resident in that population is not only stochastic, but also in an active “trade-off” against available intracellular protein stocks [46-50]. In fact, most scRNA-seq data sets exhibit gene-cell matrices that are not only characterized by their sparsity [51] but also by their near-unary structure, with almost all accrued count data having values of 1. Some argue that non-zero values in expression matrices need not be greater than 1 to yield statistical insight [52]. Mathematically, this means that in scRNA-seq datasets the values of normalized expression for most genes detected in single cells equals the inverse of each cell’s total UMI coverage. For that reason, in cases where two distinct populations exhibit similar UMI coverage per cell (equal *denominators*), differences in normalized expression of a gene based on 1-counts (equal *numerators*) may be less informative than knowing whether the gene is detected equally often in either group or not. Accordingly, in cases where UMI coverage differs substantially between populations, gene representation may be the only direct metric to ascertain whether normalized expression differences reflect either a net difference in expressed transcripts (different *numerators*), or instead, depict normalization biases deriving from mismatched UMI coverages between cell populations (different *denominators*). In the SALSA workflow we introduce metrics for gene representation rates per cluster as an important, often overlooked, parameter in scRNAseq analysis. We argue that it is critical to consider gene representation rates when analysing scRNAseq data not only because of its general sparsity, but also due to fundamental differences in what defines a DEG in “bulk-wise” vs. single-cell scales. In “bulk-wise” RNAseq, the contribution of transcripts from individual cells to a grand total within a cell conglomerate is “averaged out” and compared to those from other cell conglomerates as a continuum. In the case of sciRNAseq, individual cells either manifest countable transcripts from individual genes or not. Therefore, in scRNAseq data “average” expression differences between cell conglomerates for one gene can result from having all cells from one group expressing less transcripts than all cells in another, having transcripts scattered unevenly or in different proportions between two cell groups that express them, or a combination of both cases.

Clearly, these features of scRNA-seq data present significant challenges to performing benchtop validation. This raises a critical question: what are the best genes to test inferred cell types against given a list of candidates from scRNA-seq data? This issue of novel biomarker validation is magnified when scRNA-seq technologies are introduced to discriminate cell types within embryonic organs, novel model species, or diseased tissues that lack reliable landmark genes to validate against by alternative assays. To ameliorate these concerns, in the SALSA workflow we introduced a novel design to include additional gene strata with increasing levels of statistical stringency (Fig. 1E). Using this approach, SALSA can define a robust subset of discriminative genes insensitive to technical noise or representation rate differences between inferred cell populations. First, LSTNRs are re-tested without resolution weights across cell majors based on expression (log-fold) and representation rates combined, followed by *post hoc* pairwise comparisons between cell majors, to determine a subset of differentially expressed genes (DEGs) showing at least as many significant pairwise differences as the number of comparisons possible for each cell major. The goal of this step is not only to identify which genes among LSTNRs are significant between cell majors regardless of their position on the dynamic range of the scRNA-seq assay, but also which LSTNRs may be expressed in a mutually exclusive manner between cell majors. For example, a biomarker expressed in only 1 among 5 cell types should show at least 4 pairwise-significant comparisons, namely 1 vs. the remaining cell types. Therefore, we refer to such a minimum required number of pairwise differences suggestive of mutual exclusivity as the *cardinal number*. The resulting subset of differentially expressed genes (DEGs) is interrogated further to tease out if they have a cardinal number of pairwise-significant differences greater than SNR=1 between cell majors (Δ_Log2FC_ > 95% TI_SSR_). Genes passing this criteria are referred to as DEGs with reproducible expectation estimates (DEGREEs) because of their prospective resilience to technical noise, analogous to their bulk RNA-seq counterparts in the LSTNR method [45]. Finally, to extract candidate marker genes that are minimally impacted by representation differences between inferred cell populations, we perform unweighed log-fold expression ANOVA of DEGREEs across cell majors, and only retain those that exhibit cardinal pairwise significant differences greater than noise levels. We refer to this final subset of genes as Profilers, since they are facultative genes that lie within sequencing dynamic range based on their mean UMI coverage across single cells (SGs), statistically distinct and beyond technical noise from the average of all cells combined (LSTNRs), with statistically significant expression differences between cell types (DEGs) that are greater than noise levels and likely mutually exclusive (DEGREEs), and regardless of the proportion of cells the gene is expressed (or incidence) in each separate group. These features make Profiler genes conducive to orthogonal validation efforts by flow cytometry, ISH, or qPCR assays.

The governing principle behind SALSA is that of parsimony: the transcriptomes of cells from an organism are more alike than different, as they all use similar systems to satisfy basic needs. The exceptions arise in the form of specialized machineries encoded by specific genes. When one of such machineries is characteristic to certain cell populations, it confers those cells with the ability to fulfil unique functions within a multicellular system. SALSA shifts focus towards clarifying such distinctions by extracting sets of differentially expressed facultative genes from individual specimens, and then compiling lists of “re-incident” DEGs across multiple independent biological replicates. Next, SALSA deposits cross-replicate DEG data, regardless of scRNA-seq platforms, into a consensus expression matrix and infers cell profiles supported across biological specimens regardless of the platform used to sequence them. Inferential statistical power increases as candidate discriminant genes drop out along the SALSA workflow. Along the way data from additional independent replicates is tallied and the random effects of sequencing noise are muted, whereas “anecdotal” DEGs detected only in particular specimens or platforms lose statistical support.

From a statistical perspective, the onus that SALSA demands from facultative genes to score as consensus DEGs leads, in the end, to a narrow list of highly reproducible biomarker candidates. Unlike an all-inclusive catalogue of highly variable genes, the candidate biomarkers reported by SALSA must be mutually exclusive across identified cell types and agnostic to the choice of scRNA-seq platforms. Therefore, even though SALSA’s consensus DEGs are often smaller gene sets than those reported by alternative pipelines, they represent a minimal set of cell-specific markers with the largest *a priori* probability of success for experimental validation, and a financially viable option at that.

### SALSA identifies distinct cell types in the PBMC 3K “silver” standard dataset based on facultative genes only

#### The PBMC 3K data set exhibits a near-unary architecture

To evaluate SALSA, we analysed a publicly available “silver” standard dataset that is widely regarded for its single-cell coverage richness: the frozen Peripheral Blood Mononuclear Cells data set with 2,700 barcodes (or PBMC 3K set) available through 10X Genomics. This dataset was originally produced by 10X Genomics from a single Illumina NextSeq 500 high-output flow cell run [36]. As is, the PBMC 3K set is available in a pre-filtered fashion, in that each of the represented 2,700 barcodes is presumed to represent a single-cell. With this initial assumption at hand, we first compiled a list with all initially detected and uniquely mapped templates represented in the dataset, for a grand total of 6,390,631 barcode×UMI combinations aligned uniquely to 16,634 annotated genes in the hg19 reference genome. After tallying UMIs, we obtained a total of ∼2.3M barcode×gene data-positive count data.

Considering that a maximum allocation, or span, of 44.9M distinct count data is available in the 2,700×16,634 gene-cell matrix, it becomes clear that the “rich” PBMC 3K set is in fact a notably sparse one with the raw mappable output accounting for only ∼5.1% of the available gene-cell matrix space (Table 1). Further inspection of the data-positive fraction from the PBMC 3K gene-cell matrix revealed that ∼1.6M of all ∼2.3M data-positive count fields (∼70%) had a value of 1, followed by ∼0.3M (∼12%) and ∼0.1M (∼4%) data-positive fields with counts of 2 and 3 respectively. Combined, these three subsets (of read counts of 1-3) contained alignments for all 16,634 genes represented in the PBMC 3K library. Notably, 1-valued fields alone accounted for 16,588 (99.7%) of all aligned genes. In contrast, we found that of the remaining ∼0.32M (∼14%) of data-positive fields comprised a much smaller share of 3,929 (or 23.6% of) detected genes. These genes had 4 or more counts, with up to 419 total mappings for the single-most abundant barcode×gene combination that aligned to the FTH1 locus. We also observed that ∼0.27M (or 84%) of all multi-count data (4 or more counts) was made of alignments to only 166 (1%) of detected genes. Among the 166 “overrepresented” genes in this fraction of the PBMC 3K library we found 8 protein-coding mtDNA genes, 75 ribosomal protein subunits, 8 HLA chains, and housekeeping genes like β-actin, GAPDH, and vimentin (Table S1).

**Table 1.** Sparsity analysis of the PBMC 3K silver standard dataset by gene stratum.

| Total UMI detected: | 6,390,631 |  |  |  |
| --- | --- | --- | --- | --- |
| Gene-Cell Matrix Span (16,634×2,700): | 44,911,800 |  |  |  |
| <u>Data stratum (genexcell block size \| span)</u> | <u># Fields</u> | <u>% Matrix</u> | <u>% Stack</u> | <u>% Block Span</u> |
| **Total (16,634×2,700 \| 44,911,800)** | **2,286,884** | **5.1%** | **100%** | **5.1%** |
| constitutive genes (252×2,700 \| 680,400) | 536,804 | 1.2% | 23.5% | 78.9% |
| facultative genes (3,305×2,700 \| 8,923,500) | 1,308,249 | 2.9% | 57.2% | 14.7% |
| *SGs (2,519×2,700 \| 6,801,300)* | *1,046,003* | *2.3%* | *45.7%* | *15.4%* |
| *LSTNRs (2,519×2,700 \| 6,801,300)* | *820,191* | *1.8%* | *35.9%* | *12.1%* |
| *DEGs (1,244×2,700 \| 3,358,800)* | *558,892* | *1.2%* | *24.4%* | *16.6%* |
| *DEGREEs (464×2,700 \| 1,252,800)* | *209,238* | *0.5%* | *9.1%* | *16.7%* |
| ***Profilers (462×2,700 \| 1,247,400)*** | ***209,089*** | ***0.5%*** | ***9.1%*** | ***16.8%*** |
| rare transcripts (13,077×2,700 \| 35,307,900): | 441,831 | 1.0% | 19.3% | 1.3% |

Due to the sparsity and low-count UMI tallies in gene-cell matrices from scRNA-seq data, when developing SALSA we opted for imputation-leveraging algorithms like SVD and IRLBA that are designed to handle sparsity dynamically [44]. If missing-data fields are explicitly replaced with values of zero instead, file sizes increase dramatically, approximately ∼20-fold in the case of the PBMC 3K data set. To analyse such a dataset, large volumes of dynamic memory storage and computational processing power for data post-processing are required to queue and retain all the additional zeros that were artificially imposed on the gene-cell matrix. In practical terms, leveraging sparsity-capable algorithms to analyse scRNA-seq datasets can represent the difference between performing expression analysis for thousands of single cells using a laptop vs. having to rely on high-performance parallel processing networks to complete the same analysis.

#### PBMC 3K expression matrix focusing by a P_C_-P_D_ parametric sweep

To minimize our data footprint, we assembled the ∼2.3M data-positive barcode×gene UMI count data in a stacked format. Although this stack represents 100% of data-positive gene-cell fields, it occupies only ∼5.1% of the footprint from a traditional zero-filled gene-cell expression matrix. Next, total-per-barcode UMI counts (i.e. per-cell coverage) were recorded for calculating normalized barcode×gene expression rates in UMI-per-Thousand UMIs (UPT) units later in the workflow. Because in the 10X Genomics curated 3K PBMC dataset each barcode is regarded as a single cell, no barcode filtering by coverage was required in this instance. We tallied and recorded aggregate UMI counts per aligned gene using our P_C_-P_D_ mixture model to interrogate total-per-gene coverages (Fig. 2A). The SALSA P_C_-P_D_ mixture model was then used to determine the best candidate subset of genes ranking between low overall detection rates (i.e. rare transcripts) and extraordinarily high counts at “outlier levels” across the board (i.e. constitutive genes).

**Figure 2.**
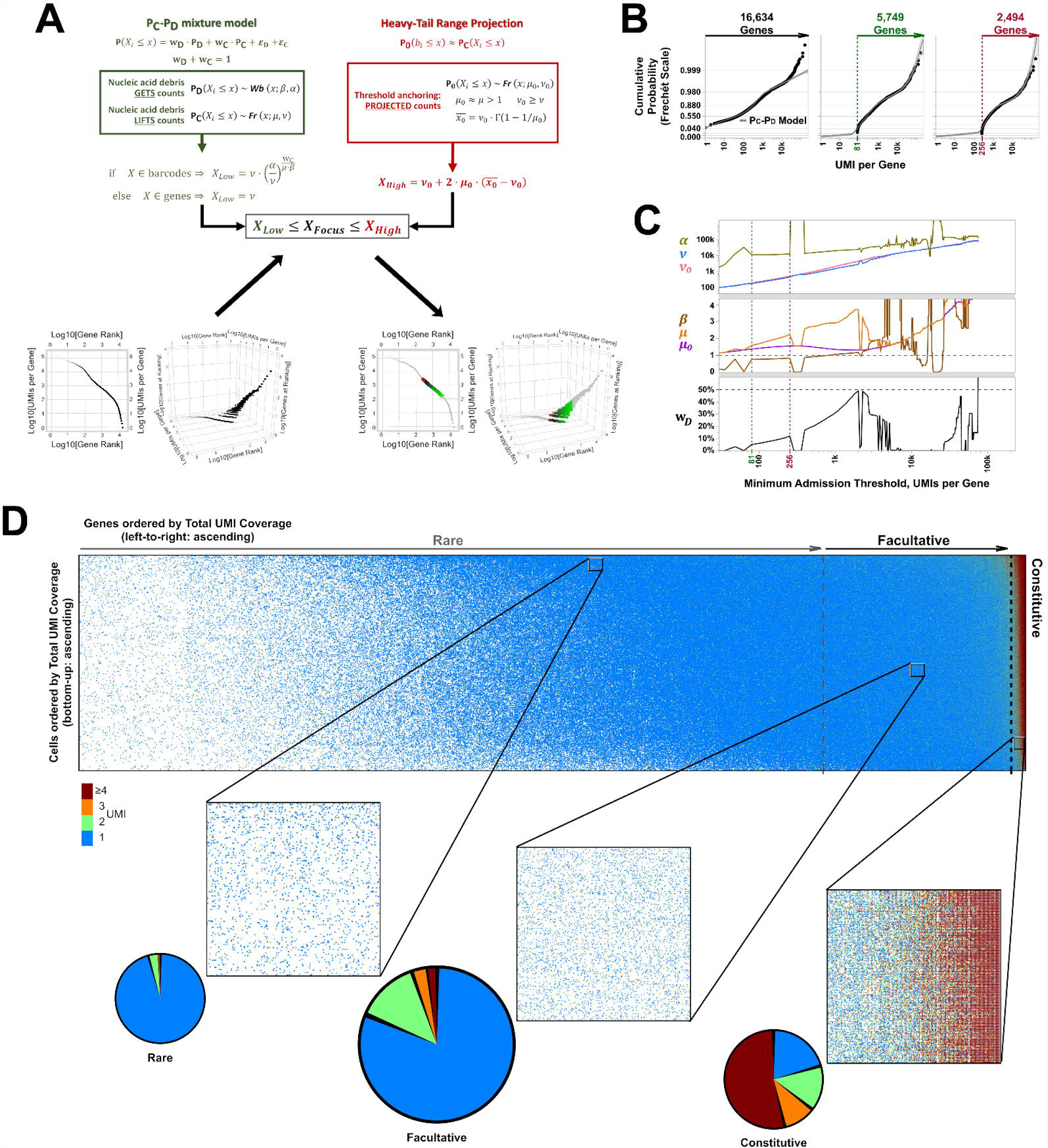
Gene-cell expression matrix focusing by SALSA on the PBMC 3K silver standard dataset. (A) Parametric focusing in the SALSA workflow. Traditional gene knee plot displays of the PBMC 3K data set with an additional z-axis showing numbers of genes sharing ranking positions (log-scale), showing all detected genes (bottom left) and highlighting inferred facultative genes (bottom right) through matrix focusing. Quantile fitting of a P_C_-P_D_ mixture model and a heavy-tailed projection model on per-gene coverages was used to estimate parametric factors and calculate “inlier” coverage bounds (top). Inferred facultative genes are shown in a low-to-high total UMI coverage color gradient (green-to-black-to-red; bottom right). (B) Example quantile plots for P_C_-P_D_ mixture model fitting at varying minimum coverage admission thresholds over all 16,634 aligned genes: no threshold (left, black label lettering), 5,749 aligned genes with >81 total UMI counts (middle, green label lettering), and 2,494 with >256 total UMI counts (right, red label lettering). (C) Parametric sweep with rising minimum coverage admission thresholds per gene of P_C_-P_D_ and heavy-tailed projection models. Green and red vertical dotted lines demarcate the span of best-fit parameters for the inferred facultative gene regime, flanked by numerical solver instabilities, that correspond to quantile plots in (B) with matching label colors. (D) Graphical representation of data sparsity in the 16,634×2,700 gene-cell expression matrix of PBMC 3K. Dots in the large rectangular frame (top) represent individual count values throughout the gene-cell expression matrix based on accrued sequencing data; missing data fields are blank. Vertical dotted gray lines demarcate the estimated boundaries between rare, facultative, and constitutively expressed genes. Make-up of count data values among data-positive fields (bottom); blow-up windows of ∼300 genes × 230 cells each for count data in the PBMC 3K expression matrix for rare, facultative, and constitutive gene regimes at high, middle, and low per-cell total UMI coverages, respectively.

To estimate ranges of “inlier” per-gene coverage scores at different minimum UMI count cut-offs, both the P_C_-P_D_ mixture model and a heavy-tailed range projection model are iterated from the bottom-up (Fig. 2B). Firstly, all detected genes with 1 or more total UMI counts are admitted for quantile regression. Best-fit parameters are then numerically solved and recorded for further analysis. To calculate this the minimum threshold of total UMI counts for gene admission is raised, and fits are repeated until the entire data stack is discarded. In our implementation, the minimum threshold is updated between iterations in squared steps (e.g. a first minimum cut-off at 1 total UMIs, second at 4, third at 9, etc.) to correspond with the 2-dimensionality of gene-cell matrices. The output from this parametric sweep consists of 7-parameter value sets from the P_C_-P_D_ mixture model (2 scale factors, 2 shape factors, and a non-zero and non-negative partial contribution rate from either the P_C_ or P_D_ function) and the heavy-tailed range projection model (1 scale factor, 1 shape factor) combined. The 5-parameter set from the P_C_-P_D_ mixture model described a best-fit distribution that contains most of the observed per-gene coverage values; the 2-parameter set from the heavy tailed projection model is used to estimate an upper bound of admissible “inlier” per-gene coverages (Fig. 2A and 2B; refer to Fig. S1 and Supplementary Materials for additional details).

Because of the nature of extreme value distributions, each fit is primarily driven by the make-up of scores closest to baseline, i.e. a minimum discriminant threshold of background vs. signal. As the threshold is raised, the estimated average and projected “inlier” maximum scores of per-gene coverages predicted by the best-fit parameters also rise (Fig. 2B). In a case where the admitted data is dominated by rare transcripts, the projected “inlier maximum coverage” is small. On the other hand, if admitted data is dominated by facultative genes, the projected maximum is larger. However, coverage data and the P_C_-P_D_ mixture model have different support (discrete vs. continuous values). This mismatch in statistical variable support can be exploited. The SALSA parametric sweep updates minimum UMI thresholds between iterations using discrete-valued squared steps, making regression algorithms susceptible to numerical singularities when estimating a continuous-valued best-fit distribution. This computational signature occurs at sudden or steep transitions in the distribution between low and high coverage values just before the minimum coverage cut-off. At such inflection points numerical solvers become unstable, leading to “spikes” in the values of best-fit P_C_-P_D_ parameters between successive iterations that would evolve smoothly otherwise. For example, “infinity” shape factors or zero-valued (machine-precision) partial contribution rates will both lead to “spikes” appearing in the chart of best-fit P_C_-P_D_ parameters (Fig. 2C). These “spikes” in the plots of P_C_-P_D_ parameter values vs. coverage cut-offs serve to highlight transitions between rare, facultative, and constitutive genes based on the ranking of their UMI coverages relative to all other detected genes. As a result, the P_C_-P_D_ parameters estimates are themselves tied to the empirical coverage data through a parametric approach. Therefore, our filtering strategy used in the SALSA workflow provides a systematic route to back-calculate ranges of total UMI coverages that distinguish facultative genes from all others represented in the data.

Based on the P_C_-P_D_ parametric sweep of the PBMC 3K data set, we partitioned the 16,634 detected genes into three categories: 13,077 rarely aligned genes (1 – 168 total UMIs each); 3,305 facultative genes (169 – 2,799 total UMIs each); and 252 constitutive genes (2,814 – 161,685 total UMIs each). The structure of the PBMC 3K data set was notable in that the data-positive fields were unevenly apportioned among the three gene coverage regimes (Table 1 and Fig. 2D). For example, rare transcripts had the largest available real-state within the gene-cell matrix, a 13,077×2,700 *block* with an allocated span of 35.3M possible count data fields. However, this real-estate added up to the smallest fraction of accrued data overall, accounting for 0.4M of the 2.3M data-positive count fields (equal to ∼19.3% of data stack). There was little improvement in block-level sparsity relative to the rare transcripts gene-cell matrix occupancy (1.3% block span vs. 1.0% matrix span); additionally, a 95.8% rate of 1-valued count fields was observed. In practical terms this meant that SALSA labelled these transcripts as *rare* because they were detected few and far between, peppered throughout the matrix at frequencies reminiscent of indiscriminate sequencing artefacts, and presumably without apportionment bias among single cells. In contrast, constitutive genes had the smallest available real-estate overall, only a 252×2,700 block, but were richer in accrued data, accounting for 0.5M data-positive fields (equal to 23.5% of stack). In addition, this real-estate was also substantially less block-sparse (78.9% block span vs. 1.2% matrix span) and dominated by multi-count data-positive fields (20.6% vs. 53.7% rates of 1-valued vs. ≥4-valued count fields, respectively). This suggests that many of the constitutive genes were often, or always, detected multiple times in most, if not all, single cells. Designation of these genes as *constitutive* is also supported by the fact that each of the 166 genes designated earlier in the workflow as “overrepresented”, based on their predominantly multi-count data make-up, were parametrically assigned under this gene stratum.

Everything considered, the 3,305×2,700 facultative gene block was the richest and compiled more accrued data than the other two blocks combined. Facultative genes harboured 1.3M data-positive fields (equal to 57.2% of stack) at higher real-estate occupancy relative to the gene-cell matrix overall (14.7% block span vs. 2.9% matrix span). Compared to the other two gene blocks, facultative genes showed an intermediate diversity in the make-up of count values with 81.3% of the data coming from 1-valued count fields and 13.5%, and 5.2% of the data making up the rates for 2-valued and ≥3-valued count fields respectively (Fig. 2D). In principle, these data features would suggest many genes in this subset were detected somewhat frequently among single cells, and often in a gradient of count values. Therefore, in practical terms, these genes are *facultative*, with their transcripts neither present or absent in all cells at once; instead, some cells express them, some do not, and some express the transcripts at rates that wax and wane.

We propose that parametric focusing of gene-cell matrices for the PBMC 3K data set, and arguably, for any scRNA-seq data sets, is a systematic curation strategy that favors retention of diverse blocks of single-cell expression data for subsequent analysis. This strategy for data curation strikes a balance between data volume, computational performance, and statistical variation. Of note, we did not perform parametric focusing at the barcode level on the PBMC 3K dataset because the source files we used only reported single-cell barcodes; even then, parametric focusing on genes alone identified gene subsets to withdraw from further analysis and greatly reduced the computational data load. In the PBMC 3K data set, this initially reduced the computational cost by well over an order of magnitude ahead of performing gene expression and single-cell clustering analyses; in effect, SALSA reduced the data to be analysed to a ∼1.3M UMI count stack vs. the original ∼45M zero-filled count matrix (Table 1). In practical terms, our parametric focusing approach to pre-processing raw scRNA-seq datasets efficiently distils the informative fraction of expression data from the prominently empty-valued matrix for further analyses.

#### Baseline benchmarking on the PBMC 3K data set with SALSA

Having honed the dataset to a facultative block containing ∼1.3M data-positive fields of gene×cell UMI counts, we then performed differential expression analysis using the SALSA workflow (Fig. 1E). Grand-mean gene expression rates were calculated using “bulked” normalized expression from all single cell barcodes combined. Log_2_-fold normalized expression differences (Log_2_FC) relative to grand-mean gene expression rates were then calculated within single-cell barcodes and used for subsequent inferential tests.

Even before expression differences between individual barcodes can be inspected, there are two critical properties of expression measurements that must be demarcated first to produce a reliable statistical analysis: the dynamic range of measurement sensitivity, and the precision of expression measurements. To estimate a useful dynamic range of detectable gene expression over library background, we characterized the distribution of log-transformed grand-mean gene expression rates. We found that a thresholded normal distribution was the best-fit probabilistic model among probability functions of the exponential family. This parametric fit, which is amenable to canonical GLM-based linearization, helped estimate linear predictor values of gene×cell expression rates via GLM [45].

For a scRNA-seq library containing different cell populations, precision of normalized expression measurements is the variation from single cells relative to the average of the populations they belong to. However, in scRNA-seq libraries like PBMC 3K, the underlying populations are unknown at first. To address this issue, we used linear predictors of gene×cell expression rates to infer an initial single-cell breakdown into 7 prospective clusters by IRLBA-wrangled unsupervised hierarchical clustering (Ward’s method) [44]. Then, linear relationships between Log_2_FC scores per gene of single cells around the means of their clusters were estimated by linear regression (*R*^2^ = 0.84). The resulting sums of squared residuals (SSR) were used to define precision baselines, namely a 5%-level practical effect size δ_Log2_FC = 0.05×(6σ_Gene SSR_), and a global Log_2_FC measurement error of ±1.07 (|Log_2_FC| = 95%TI_Bulk SSR_) as SNR = 1 benchmark [45].

From the facultative block (1.3M data-positive fields, 57.2% of stack) we identified ∼76% of genes as SGs (2,519 of the 3,305 genes; Table 1 and Fig. 3A). These genes accounted for the subset of facultative genes that, once resolution and within-cluster representation rates were accounted for in combination, showed statistically significant variation among the 7 prospective clusters (Fig. 1E). As a subset of the facultative block, the SG stratum (2,519×2,700 block) harboured less data-positive fields (1.0M, 45.7% of stack) and yet, as we anticipated, the SG subset showed higher data occupancy (15.4% block span) compared to the facultative block altogether (14.7% block span). Next, we found that, of the 1.0M count data in the SG stratum, 0.82M qualified as LSTNR block data (35.9% of stack), as it cleared both the 5% practical effect sizes per gene and the global SNR thresholds based on gene×cell Log_2_FC values. Interestingly, this LSTNR stratum contained each of the single-cells and genes found in the SG stratum before effect size and measurement error thresholding. Therefore, even though SALSA is extracting an increasingly robust data subset when moving from SGs to LSTNRs, a classification which demands more statistical stringency, it did so without barcode or gene dropouts (Table 1). Conversely, since the resulting LSTNR block was of the same size as its SG predecessor (2,519×2,700), our findings also implied that filtering for gene×cell count data “above precision baseline” meant some data-positive fields were cleared out, which lowered effective data occupancy from 15.4% to 12.1% block spans between SG and LSTNR strata, respectively (Table 1). This resulted in a net 78.4% SG-to-LSTNR data retention rate for the PBMC 3K data set (Table 1).

**Figure 3.**
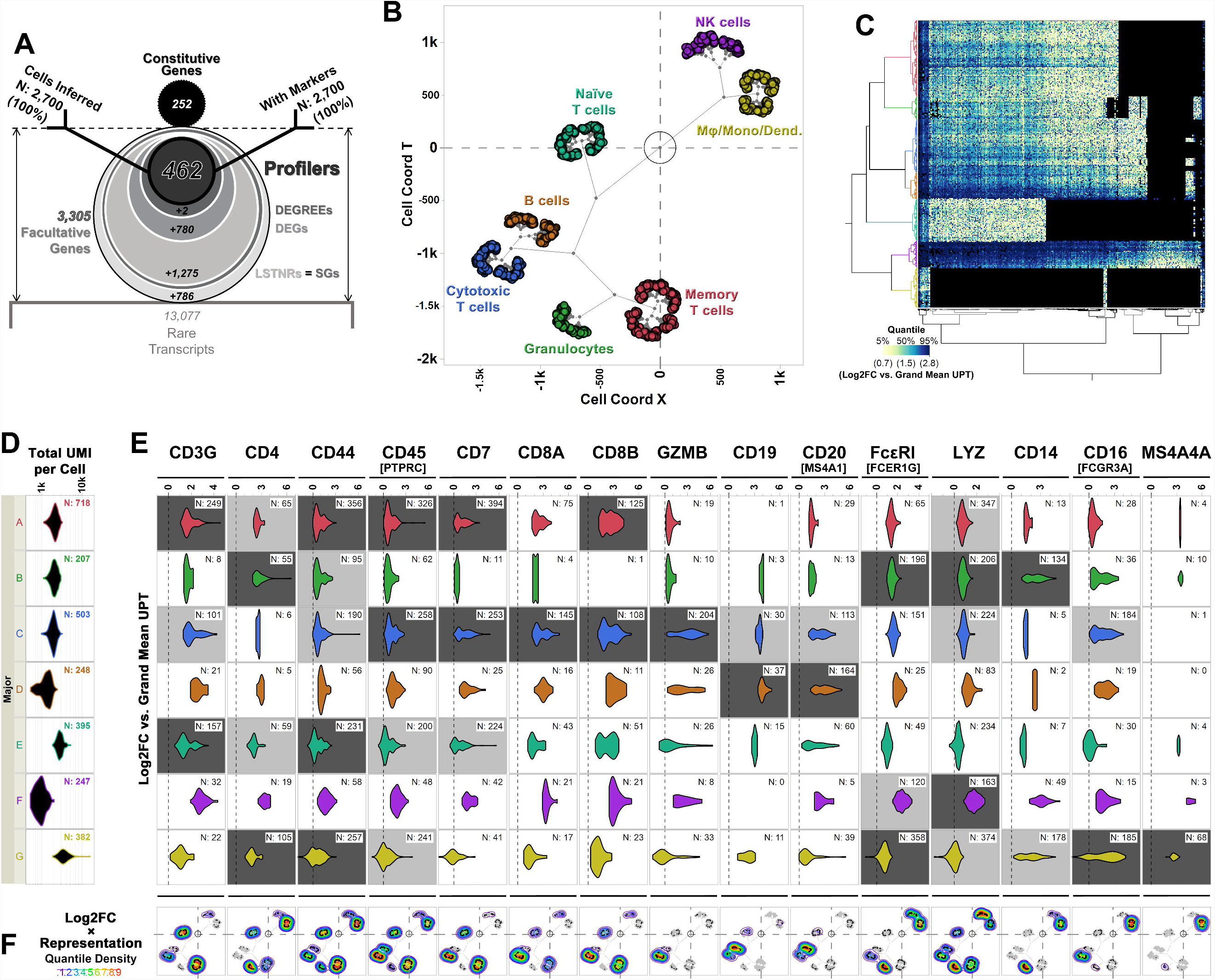
Differential expression analysis and cell type inferences in the PBMC 3K dataset using SALSA. (A) Frosty plot of gene stratification across rising levels of statistical significance in PBMC 3K. Head (black circle, top), body (largest encasing circle, middle), and base (dim gray rectangle, bottom) depict the make-up of detected genes from a single-cell library based on their constitutive, facultative, or rarely expressed status; number of facultative genes admitted past significance criteria in each stratum are also shown as encasing circles with varying sizes and grayscale intensities. Stick arms flag the gene stratum chosen as the agnostic expression marker gene set for final inferential clustering of single-cell barcodes into cell majors. Retention rates of input and output single-cell barcodes following gene stratification are represented by the relative heights of stick arms going from 100% of inferred cells with facultative gene data (left arm) to a subset of inferred cells expressing agnostic markers (right arm; e.g. 100% single-cell barcodes at the Profiler gene stratum in PBMC 3K). (B) Putative cell types matched to cell majors and their inferred transcriptional proximities displayed in latent 2D space by unsupervised clustering of mean linear predictor estimates «***B***(**θ**) » for expression rates of agnostic markers. (C) Heatmap overlay onto two-way clustering dendrograms from (B) showing increasing quantile scores of Log_2_FC values relative to library-wise UPT grand mean (tan-to-cyan-to-blue); missing data fields are shown in black. (D) Violin plots for total UMI coverage per barcode (x-axis) within cell majors; inset legends report total number of barcodes per cell major. (E) Violin plots of Log_2_FC values relative to library-wise UPT grand means (x-axis) for 15 landmark expression genes of blood cell types across cell majors in PBMC 3K; inset legends report total number of barcodes with UMIs for each landmark gene. Relative representation rates of landmark genes, i.e. the ratio of expressing vs. total cells within cell majors, are illustrated by coloring of violin plot backgrounds: in gray for all clusters with the highest Log2FC expression levels for a given landmark gene, and in a darker hue for those also showing the highest representation rates across cell majors. (F) Topographs showing the patterns of expressed landmark gene enrichment across the latent 2D space map from (B), overlayed with a non-parametric quantile heatmap for “weighed gene expression” scores, i.e. the composite score of single-cell Log_2_FC values and within-major representation rates per gene; individual expressing cells are shown as black dots throughout the 2D clustering map.

In our view, the progression of admitted data in our SG-to-LSTNR transition posits an important cautionary tale: without a proactive precision benchmarking strategy, one risks inferring cell clusters based on gene-cell expression matrices unduly burdened with measurement noise. In the PBMC 3K data set this means that with a 78.4% retention rate, the LSTNR block from the PBMC 3K data set contains the exact same single cells and barcodes as the SG block, yet it is equivalent to discarding roughly 1 out of 5 data points worth of noise before further analysis (Table 1). Both the SG and LSTNR blocks represent minute fractions of the original expression matrix, with a 2.3% and 1.8% matrix span respectively. Additionally, most of the expression matrix real-estate is earmarked for overwhelmingly sparse or 1-valued count data from rare transcripts or artefacts, accounting for ∼78.6% of the matrix allowance (Fig. 2B). In conclusion, in analysing this “silver” dataset we can clearly see how inefficient and misleading scRNA-seq data wrangling can become without performing statistical efforts devoted to either matrix focusing or precision baseline benchmarking prior to clustering of expression patterns.

#### Stratification of LSTNRs and single-cell clustering in the PBMC 3K data set

The precision-benchmarked subset of 2,419 LSTNR genes represents a diverse catalogue of candidate discriminants to distinguish cells belonging different clusters by their transcriptional signatures. Still, at this stage, these cell groupings are prospective and relatively unrefined: considering that LSTNRs represents roughly 3 out of every 4 facultative genes in the PBMC 3K data, one could counter that the initial single-cell clustering, which included non-LSTNR data, could be disproportionately influenced by data-positive fields below their own *a priori* SNR=1 estimated threshold. To address these concerns, we gathered LSTNR block data and, using the SALSA workflow (Fig. 1E) inferred 7 “cell major” groupings via IRLBA re-clustering of barcodes based on the GLM-projected expression rates, then scored representation rates per LSTNR gene within cell majors (Fig. 1E). In this context representation rates reflected the proportion of cells with detected UMIs aligning to LSTNRs.

At this point, we reached a cross-road. On the one hand, we extracted a precision-curated catalogue of LSTNR genes, whose statistical variation could be exploited to sort out underlying cell types from a corpus of single-cell transcriptomes through bioinformatic means. On the other hand, in understanding the biology behind the system of interest, it is of little advantage to have over 2,400 LSTNR genes if there is no insight on which the best candidates are to validate bioinformatically inferred cell types using on-the-bench assays like qPCR. Similar concerns can be raised regarding representation weights in SALSA. Utilising representation weights, we successfully enhanced our ability to discern between cell majors, as they account for “yes/no” expression status of genes depending on the single-cell cluster. In fact, when interrogating expression matrices dominated by 1-valued gene×cell count data, representation weights provide perhaps the only differential metric at hand that truly discriminates between inferred cell types. Yet, representation rates at the transcriptional level are notoriously difficult to validate in a quantitative fashion on the bench.

We sought to address this data-to-bench conundrum in the SALSA workflow by inspecting additional layers of gene stratification within LSTNRs, based on how prospectively can LSTNR subsets reproduce differential expression patterns in experimental assays. We convey gene stratification results hereafter using a short-hand graphical aid, the “frosty” plot, that illustrates the transition in data retention as detected genes are sifted through increasingly stringent filters of statistical significance. The frosty plot for the PBMC 3K data using SALSA-based gene stratification, and details on its assembly, are shown in Fig. 3A.

As mentioned earlier, the list of LSTNRs is iteratively focused by progressively stringent stratification of statistical significance and *post hoc* cardinality of differential Log_2_FC-based expression between cell majors (Fig. 1E). By sequentially “stripping” statistical comparisons of resolution weights, followed by representation weights, while demanding mutually exclusive *post hoc* expression differences between cell majors with statistical and above-noise significance throughout, we narrowed the pool of genes. From a 2,419-member list of LSTNR genes from the PBMC 3K data set we sifted the pool down to: a) 1,244 DEGs whose Log_2_FC pairwise differences between cell majors are statistically significant and mutually exclusive (DEGs); of which b) 464 showed differences greater than the SNR=1 noise benchmark (DEGREEs); including c) 462 exhibiting statistically distinct and mutually exclusive differences that are insensitive to stochastic incidence of transcripts, or representation, among cell types (Profilers). Notably, even though the number of retained gene×cell count data fields dropped as the number of genes decreased between strata (Table 1) each of the 2,700 single-cell barcodes in the dataset expressed between 3 – 175 of the 462 Profiler genes, with total Profiler-aligned coverage per cell ranging from 4 – 289 UMIs. Our stratification approach led to a substantial improvement on information maximization rates: the 462×2,700 Profiler block represents ∼2.8% of the gene-cell matrix allowance, and a minute portion of the accrued data overall (9.1% of the accrued data stack, equal to 0.5% matrix span). This portion accounts for the single least sparse stratum within facultative gene data (16.8% block span). Incidentally, this span is over 3× more populated as a subset than the gene-cell expression matrix overall (5.1% matrix span), as outlined in Table 1.

One underlying assumption of this approach is that LSTNR-based single-cell clustering is reasonably conserved across the stratification process. Thus, to ascertain whether that implied fidelity held true, data from the Profiler block was used in IRLBA re-clustering. Here, cell majors at the LSTNR and Profiler strata were compared to one another in terms of membership by cluster-to-cluster concordance, and hierarchical sorting by single-cell ordering within clustering dendrogram. In all, we found that single cells were largely classified into matching clusters before and after LSTNR stratification (contingency *R*^2^ = 0.91, Pearson χ^2^ *p*<1×10^−4^). These results suggest that facultative gene stratification retains underlying transcriptional profiles of single cells, thereby pointing to SALSA successfully extracting a parsimonious subset of testable, agnostically defined candidate biomarkers.

After performing single-cell clustering at the Profiler-stratum level, we first paid attention to the relative density of data in each of the 7 cell majors, originally labelled A through G. Several intricacies became immediately apparent – for example, not only were the number of cells different among cell majors, but also the distribution of total UMI coverages in each was not even-handed. Based on their clustering dendrogram (Fig. 3B and C), all 7 majors that we defined were grossly derived from three core stems: A through D represented one extreme, F and G were situated opposite, and E constituted a lone-standing intermediate stem. In the first stem, only majors A through C, which accounted for slightly over half of all inferred cells, showed similar total UMI coverage distributions among them (roughly 1,000 – 3,000 total UMI per cell; Fig. 3D). Log_2_FC expression levels for A through C were in the low-to-mid quantiles among accrued profiler-stratum count data overall (about 0.7 – 1.5 log-fold relative to the bulk-wise average; Fig. 3C). Meanwhile, major D was substantially skewed towards lower total UMI coverages, with about 500 – 2,500 total UMI per cell (Fig. 3D), and mid-to-high Log_2_FC expression quantiles (up to 2.8 log-fold vs. bulk). Interestingly, based on net Log_2_FC values, normalized expression rates in major D were about twice as high as those from the A-to-C ensemble. This is seemingly consistent with the mismatch between both ensembles in total UMI coverage, and to which count data each are normalized against. This observation implies that traditional normalized expression scores at the gene×cell level, such as UPT rates, track with total count coverage rather than intensity of expression. Accordingly, cell major E in the intermediate branch showed lower expression values relative to the A-C ensemble in the first stem by about a factor of 2. This was consistent with cells in major E carrying roughly twice as many total UMIs the A-C ensemble (about 2,000 – 5,000 total UMI per cell; Fig. 3D). Put simply, these data demonstrate that single-cell expression metrics by normalized-per-coverage rates alone are misleading.

Using IRLBA-based single-cell clustering within the SALSA workflow illustrates how interpretation of expression matrices can go awry if based solely on expression scoring without account for representation or precision benchmarking. For the PBMC 3K dataset, these risks are most evident in the F-G stem: based on hierarchical clustering (Fig. 3B, C), majors F and G are more closely related to each other than to any other cell majors, yet their total UMI coverages are at opposite extremes (500 – 2,000 and 2,000 – 16,000 total UMI per cell in F and G, respectively; Fig. 3D). In a seeming contradiction, data in major F aligns to the most Profiler genes while, at the same time, exhibits the lowest per-cell coverage among all majors; major G data make-up for Profiler genes is exactly the opposite. One explanation for such extreme behaviours may be that per-cell coverages for majors F and G are disproportionately laden with UMIs from constitutive genes, all of which are omitted from clustering tests – suggesting that either the translational (e.g. transcripts for ribosomal subunits) or metabolic (e.g. mtDNA-derived and respirasome-encoding transcripts) states of cells in majors F and G are at odds relative to each other. Another explanation could come from the effect of total coverage itself on gene stratification: for single cells whose data is dominated by 1-valued counts per gene, more total UMIs will result in lower UPT scores and a larger proportion of “muted” gene×cell Log_2_FC values below the SNR=1 threshold, which then would be excluded from the expression matrix at the LSTNR stage.

#### Peripheral blood cell types in the PBMC 3K data set inferred using SALSA

To examine whether Profiler-based unsupervised clustering tracked with transcriptional signatures from peripheral blood subpopulations, we focused on expression data from a reference subset of 15 “landmark” genes encoding 14 widely recognized protein markers (Figs. 3C, E and F; Table 2). To do so, we inspected within-cluster distributions of Log_2_FC expression levels relative to the whole-specimen collective. We accounted for “yes/no” representation rates within each cluster for landmark genes; by coloring the background of violin plots grey we highlighted clusters with the highest overall Log2FC expression levels in any given gene and showed the highest representation rates in combination (Fig. 3E). To visualise the clustering data in 2D we devised “topographs” using 2D clustering maps overlayed with a non-parametric quantile heatmap highlighting “weighed gene expression” scores. These scores represent a composite of single-cell Log_2_FC values and major-wise representation rate, defined as the proportion of positive cells per major, for a gene of interest. We use these topographs to simultaneously highlight differences in the intensity and predominance of expressed genes among clusters of like cells (Fig. 3F).

**Table 2.** Landmark genes, their gene stratum classification by SALSA, and their expression levels among cell types in the PBMC 3K silver standard dataset.

| Landmark Gene Name<br>[Entrez Symbols] | Gene<br>Stratum | Naïve<br>T-cell | Memory<br>T-cell | Cytotoxic<br>T-cell | B-cell | NK cell | Granulocytes | Monocyte &<br>M-derivatives |
| --- | --- | --- | --- | --- | --- | --- | --- | --- |
| **CD3**<br>[CD3D/E/G] | DEG | ++ | ++ | + |  |  |  |  |
| **CD4** | Facultative | + | + |  |  |  | ++ | ++ |
| **CD44** | Profiler | + | ++ | + |  |  | + | ++ |
| **CD45**<sub>RA/B/C/O</sub><br>[PTPRC] | Profiler | + | ++ | ++ |  |  |  | + |
| **CD7** | Profiler | + | ++ | ++ |  |  |  |  |
| **CD8**<br>[CD8A/B] | LSTNR |  | + | ++ |  |  |  |  |
| **GZMB** | Facultative |  |  | ++ |  |  |  |  |
| **CD19** | Rare |  |  |  | ++ |  |  |  |
| **CD20**<br>[MS4A1] | Facultative |  |  |  | ++ |  |  |  |
| **FcεRI**<br>[FCER1A/G, MS4A2] | Constitutive |  |  |  |  | + | ++ | ++ |
| **LYZ** | Constitutive |  | + | + |  | ++ | ++ | + |
| **CD14** | LSTNR |  |  |  |  |  | ++ | ++ |
| **CD16**<br>[FCGR3A] | LSTNR |  |  | ++ |  |  |  | ++ |
| **MS4A4A** | Rare |  |  |  |  |  |  | ++ |

Single-cell clustering based on Profiler genes in the 3K PBMC dataset revealed 7 distinctive single-cell clusters (Fig. 3B and C) split into two overarching transcriptome categories; the first category containing the A-D ensemble and the intermediate E stem (2,071 cells combined), and the second containing the F-G stem (629 cells combined). Within the first transcriptome category, majors A, C and E showed expression of CD3 subunits which is characteristic of T cells, whereas major D was distinguished by rich CD19+/CD20(MS4A1)+ expression consistent with a B cell phenotype (Fig. 3E) [53-59]. In addition, majors A, C, D and E all expressed CD124(IL4R), a marker shared by both T and B lymphocytes [60,61] (Fig. S2). Together, lymphoid-derived T cells (majors A, C and E) and B cells (major D) accounted for 1,864 cells; this contribution is consistent with the reported 4:1 ratio, for cells of lymphoid vs. myeloid origin in the source PBMC stock [62]. In contrast, cell major B, which represents the remaining cells from the first transcriptome category, showed a different pattern of gene expression. One of the richest CD4+ subpopulations in the entire dataset, based on its Profiler genes cohort, cell major B, was transcriptionally similar to other lymphoid-derived subpopulations it clustered with (Fig. 3C). However, cell major B was otherwise discordant when compared to its companions, majors A, C, D and E, based on landmark genes (Fig. 3E). Cells in major B had low CD16 expression, were rich in expression of landmark genes CD14, LYZ, and FCεRI(FCER1G), exhibited a myeloid-linked CD33+/CD36+ profile, and expressed the Fc-γ receptors CD64(FCGR1A/B), CD32(FCGR2A), and FcRn(FCGRT) (Fig. 3E and S1). Based on the overall pattern of landmark gene expression and other supplemental markers, we determined that the transcriptional profile of cells in major B was consistent with granulocytes [54,63,64].

To further split out the functions of majors A, C and E we exploited the unique characteristics of lymphoid-derived T cells subclasses. Transcriptionally, lymphoid-derived T cells exhibit differences based on their maturation state, regulatory vs. inflammatory roles, and degrees of antigen-presenting and antigen-discovery competence [53,55,57-59,61,65-68]. However, transcriptional differences between majors A, C and E in the PBMC 3K data set (depicted using violin plots; Fig. 3E) were too subtle to be matched to discrete T cell subtypes based on normalized expression rates alone. In this context, inspection of gene representation rates proved critical in assigning lineage identity to cell majors; here topograph visualisation outperforms violin plots by simultaneously revealing log-fold expression, representation rates, and the location of expressing individual cells in clustering maps for a gene of interest (Fig. 3F). Using topographs we observed gradients in CD3+ expression consistent with known transcriptional changes during T cell maturation across cell majors A, C and E. In major E we revealed a CD3^hi^/CD4+/CD44^hi^/CD45+/CD7+ naïve T cell signature, and then isolated two related phenotypes: memory T cells enriched in major A (CD3^hi^/CD4+/CD44^hi^/CD45^hi^/CD7^hi^), and cytotoxic T cells in major C (CD3+/CD4-/CD44+/CD45^hi^/CD7^hi^/CD8^hi^; Fig. 3E and F). In summary, using SALSA we identified majors E, A and C as naïve, memory and cytotoxic T cells respectively; this is in agreement with varying degrees of enrichment for additional T cell maturation markers, including CCR7, CD27, IL2R (CD25) subunits, and CD127(IL7R) (Fig. S2) [56,65,67,69,70]. Similarly, our identification of major D as the B cell subpopulation agreed with the near exclusive expression of additional B cell population markers, such as CD23(FCER2), CD32B(FCGR2B), and CD79 subunits (Fig. S2) [56,71].

Arrangement of cell subpopulations within the first transcriptome category, encompassing majors A, C, D and E, revealed that SALSA-based unsupervised clustering can successfully reconstruct matches between cell types and their coordinated biological functions. For example, we found that majors C and D had the closest transcriptomic similarity among all clustered cell subpopulations (Fig. 3B). Notably majors C and D correspond to cytotoxic T cell and B cell lineages which are mature subpopulations with closely linked and converging physiological roles [57,58,60,61,71]. Likewise, the inferred proximity between memory T cells in major A, and granulocytes in major B in the clustering map (Fig. 3B) suggests a significant interaction between those two cell types. Physiologically this relationship is widely recognized; subsets of granulocytes are known to perform antigen-presenting roles that help coordinate CD4+ T cell activation [54,63,64].

The second transcriptome category constituted majors F and G. This was puzzling, not only because majors F and G had strikingly different total UMI coverages, as mentioned earlier, but also because we found that expression of blood cell-type markers in major G cells had the strongest similarities with granulocytes (major B) in the first transcriptome category. Both majors G and B shared similar representation rates for landmark genes CD4, CD14, and FCεRI(FCER1G) (Fig. 3E), carried a CD33+/CD36+ myeloid signature (Fig. S2), and had matching logistic regression trends for Profiler genes responsive to microbial infections, such as AP1S2, CNPY3, FPR1 and RGS2 [72-75] (Fig. S3). However, major G cells clustered apart from granulocytes in major B and were also distinct in critical ways; for example, major G cells showed much higher expression levels of monocytic and macrophage-enriched genes CD16(FCGR3A) and MS4A4A, respectively (Fig. 3B, C and E) [56,76,77]. We concluded that cell major G represents a combined pool of monocyte-derived subtypes including monocytes, macrophages, and mono-derived dendritic cells. This conclusion was based on two key factors: an implied myeloid origin, like that of antigen-presenting granulocytes in major B, and transcriptional enrichment for genes involved in pathogen recognition processes that are characteristic of leukocytes from the innate immune system [54,56,66,73,74,76-78].

Given this backdrop, we surmised that if clustering proximity in the second transcriptome category evokes converging physiologies between non-myeloid cells from major F and myeloid-derived cells from major G, irrespective of their strikingly different total UMI counts, then majors F and G should both present key ontologies. To test this, we asked whether majors F and G shared a minimal set of abundantly detected Profiler genes which: a) were absent from all other cell majors; and b) significantly enriched for critical innate immunity processes or molecular functions. After surveying weighed expression rates by multinomial logistic regression, we found that both majors F and G prominently expressed the Profiler genes HIST1H2AC, PF4, PPBP, and SDPR(CAVIN2). These genes all participate in anti-parasitic NETosis processes, mediate leukocyte chemoattraction and degranulation, and are regulated by calcium-sensitive protein kinase C activity [79-85]. Importantly, most other cells in the PBMC 3K dataset did not express any of these Profiler genes or exhibit their associated ontologies (Fig. S3).

Still, the transcriptomic profiles of cells in major F were evidently distinct from monocytic cell types, not only by their coverage, but also because of critical landmark gene signatures. The genes FCεRI(FCER1G) and LYZ were both detected in almost 2 out of every 3 cells from major F, which expressed them at the richest Log_2_FC levels among all cell majors relative to the bulk mean. Both these landmark genes showed their lowest Log_2_FC levels across the dataset among cells from major G, even though transcripts from both FCεRI(FCER1G) and LYZ were detected in almost every single major G cell. In contrast, CD16 expression was all but missing from major F, and most prevalent in major G cells overall. Therefore, we used four defining features to derive the identity of the cells in major F: 1) their strong ontological relationship with monocytic cell types, 2) their relative frequency in the data set of ∼9% of the total (247 single cells), 3) their non-myeloid origin, and finally, 4) their inferred regulatory functions linked to innate immunity within PBMC fractions. From these features we deduced that cells in major F correspond to lymphoid-derived natural killer (NK) cells [62,86-91].

Finally, it is worth mentioning that segregation of T cell subtypes, B cells, and antigen-presenting granulocytes under the same transcriptome category when using the SALSA workflow was consistent with an underlying and powerful biological feature: those four cell types constitute the adaptive immune system. Conversely, the second transcriptome category depicted the main players in the innate immune response: NK cells and monocyte-derived cells. We found that unsupervised clustering of single cells inferred by SALSA recapitulated the expression patterns of traditional marker genes and proportions of cell types expected from PBMC specimens without data preconditioning. Thus, by performing single-cell profiling on the “silver” standard PBMC 3K dataset, we demonstrate the core strengths of the SALSA workflow. With SALSA, a minimal fraction of well-resolved expression data from agnostic Profiler genes successfully sorted like cells, recapitulated experimentally demonstrated transcriptional signatures, and retained latent linkages that evoke converging physiologies among interconnected cell types.

#### SALSA extracts lineage-specific transcriptional signatures while circumventing fleeting retention and high-dropout rates of mRNA templates encoding protein-based markers

Regardless of technique, scRNA-seq data registration, the ability to trace provenance of sequenced data back to each of the starting cells, requires that all templates in the cDNA pool from a cell carry a shared tell-tale barcode sequence. This sequence should be discernible from amplified dsDNA templates coming from elsewhere, such as nucleic acid debris, PCR concatemers, or other cells. With few exceptions, scRNA-seq library preparation chemistries for high-throughput scRNA-seq platforms use anchored oligo dT sequences hybridized to 3’ polyadenylated tails of mRNA molecules and then prime cDNA synthesis. Reverse transcription into cDNA is coupled with cell-level and molecule-level barcodes to trace reads back to initial mRNA templates, and combined with universal priming sequences, termed “handles”, that permit multiplexed amplification of single-cell cDNA banks. From that point forward, the raw sequencing reads generated are deplexed into cell and template of origin, and aligned to a reference genome [92].

Off the shelf instruments for state-of-the-art scRNA-seq technologies still predominately utilise short-read sequencing-by-synthesis (SBS) platforms that use reversible terminator-bound dNTPs protocols because of their higher output in terms of read counts per assay. For short-read scRNA-seq, barcoded cDNA templates are sheared within the 300-700 bp size range. The resulting sheared cDNA pool includes end-tagged fragments, spanning between the appended “outer” barcodes and “inner” cleaved exomic ends. These end-tagged cDNA fragments are then enriched by targeted amplification from their appended universal handles. The final end-tagged scRNA-seq library is used in bridged paired-end sequencing, surveying “outer” barcode ends first, and “inner” exomic ends second. The paired reads are deplexed by their barcodes in first reads and aligned by their exome sequences in second reads bioinformatically.

In the most commonly used chemistries, universal handles and barcodes are fused to poly-A^+^ insertion sites of mRNA molecules. In this context, bridged paired-end sequencing allows each read from sequenced fragments to serve different purposes. By splitting the roles for deplexing and alignment between the end of sequenced fragments this approach aims to circumvent mappability issues from oligo dT mispriming. The dT regions can anneal randomly to points inside poly-A^+^ tracks in RNA templates, but also to degraded gDNA debris. When oligo dTs that anneal with mRNA molecules fail to anchor by stop codons, single-end sequencing of their end-tagged cDNA copies yields unmappable A/T run-ons. These poly-A^+^ run-ons cannot be skipped in single-end sequencing when using reversible terminator chemistries because each base is called one at a time. However, with paired-ending, library templates are flipped once “outer” barcode sequences have been collected, then sequencing is resumed from the opposite ends outwards. This strategy is more likely to capture mappable information from the “inner” ends of tagged cDNA fragments.

After trimming off the barcode and sequencing adapter from the sequenced reads, paired-end short-read sequencing usually yields deplexed “inner” data aligned within <500-bp from the cistromic 3’ end of mRNA molecules. In that sense, paired-end short-read sequencing of scRNA-seq libraries does without full exome representation. Based on GENCODE annotations, aligning only 500-bp 3’-end mRNA fragments allows sampling the human exome with ∼18.0 Mb total information, or 2.7-fold less nucleotides than full-transcript sequencing (∼47.9-Mb length, 1,350-bp full-transcript size average, ∼36,000 annotated genes) [93],. Furthermore, based on an average ∼95% unique mappability rate with ≤2 nt mismatches for the protein-coding exome using 100-kmers [94], the total mappable reads required for unambiguous sampling of 3’-ended human exomes can be reduced an additional 5-fold. Overall, these estimates project a mappable 3’-end sampled exome of ∼3.8 Mb at 1× coverage, roughly 13 times less sequencing reads per library than needed for full exome representation by traditional RNA-seq [94].

Still, using short-read 3’ end-tagging chemistries in hopes of increasing gene call rates in scRNA-seq libraries is not without compromises from a benchtop perspective. It is critical to remember that the bulk of RNA molecules in any given cell consists of different types of RNA species; over 90% of those RNA templates are derived from rRNA species that can harbor poly-A^+^ tracks, with only about 5% - 10% of total RNA representing mature mRNA transcripts [95-99]. Additionally, although numbers of native RNA molecules vary considerably among individual cells, the rates of RNA capture during single-cell cDNA synthesis, and the fractions of those represented in scRNA-seq libraries, are limiting [92]. All things being equal, other significant bioinformatics challenges are to be expected when aligning 3’ end-tagged fragments generated in scRNA-seq libraries to reference genomes. A short-read “targeted” sc-RNAseq exome spanning ∼500-bp cistromic windows is at odds with the architecture of 3’ UTRs in eukaryotes, which are often larger in size. In humans, for example, short 3’ UTRs predominate during development; these are spliced at proximal poly-A^+^ sites and result in median 3’ UTR lengths of ∼400 bp. Alternative 3’ UTR isoforms found in differentiated tissues typically are in the range of kilobases [100]. These features make it clear that data from typical 3’-end scRNA-seq libraries, based on 300-700 bp size fragments, disproportionately relies on uniquely mappable segments from within 3’ UTRs, and not on sequences from within protein-coding exons. In other instances, anchored oligo dT primers used in reverse transcription may hybridize away from stop codons of mRNA templates and within their poly-A^+^ tails; with ≥300 bp average lengths for mammalian mRNA poly-A^+^ tails [101], 3’-end fragments from unanchored cDNA templates often yield sequenced reads consisting of unmappable A/T run-ons.

Considering all these factors, it is somewhat unsurprising that genes encoding destabilizing *cis*-regulatory elements in 3’ UTRs would be harder to detect by scRNA-seq, as high mRNA turnover is at odds with maintaining high steady-state mRNA levels necessary to capture them for cDNA synthesis [102]. Less appreciated is that since 3’ UTRs encode evolutionarily conserved binding sequences and repetitive elements modulating post-transcriptional regulation of mRNA processing, one may find that sequenced reads from 3’ UTRs of some key developmental regulators may not be uniquely mappable. If this is the case, lineage markers and stress-responsive genes may be particularly susceptible to both low mRNA capture and high sequenced read dropout rates, leading to poor total UMI counts in 3’-end scRNA-seq libraries. An example of this concept would be genes regulating stem cell transitions into mature cell types. In the PBMC 3K dataset, we see this concept illustrated clearly in two critical markers: the membrane-bound NK cell marker CD56, and the Treg/Tsup-specific nuclear transcription factor FOXP3.

Expression of cell surface molecule CD56, formally known as neural cell adhesion molecule (or NCAM), is the most reliable and widely used epitope-based marker for sorting NK cells by flow cytometry [56,66,90]. For decades, NK cell subtypes have been defined in relation to their intensity of CD56 protein expression among blood cells. Regulatory NK cells in peripheral blood are CD16-negative and “bright” for CD56, whereas inflammatory NK cells are defined as CD56^dim^/CD16+ and tissue-resident. All other leukocytes and hemopoietic cell types are CD56-negative, including Lin-cells (CD14-/CD3-/CD19-/CD16-/CD56-) which are rarely detected in peripheral blood [54,56,66,86,89-91]. Despite CD56 being a *bona fide* epitope marker, the PBMC data set only contained 14 total UMIs aligned to the CD56 gene (14 individual cells, 1 UMI each). This by itself was far too little information to infer an NK cell cluster with; instead, we recognized NK cells based on other features in combination. We believe the disconnect between expected NK cell numbers in the specimen detected by alternative markers [62] and the poor counts for reads uniquely aligned to the CD56 locus reflect mappability challenges for the CD56 3’ UTR sequence. According to the hg19 annotation (100% homologous to its hg38 version), the CD56(NCAM1) gene encodes a 3’ UTR with three poly-A^+^ insertion sites, with the most proximal sites located at 1.2 kb, 1.1 kb, and 0.5 kb downstream from the mRNA stop codon. Additionally, the CD56(NCAM1) gene contains a 3’ UTR (CA)_n_ single-repeat, and a 3’ UTR (TTTC)_n_ simple-repeat (Fig. 4A). Among reported spliced ESTs, only the (CA)_n_ element is retained and is located 200 bp within the 3’ end of DA492761, which is the longest spliced EST overlapping the genomic annotation of the NCAM1 3’ UTR. With the limited information on CD56 mRNA stability in NK cells, we surmise that low mappability of simple-repeat elements prevented unambiguous alignment of CD56 end-tagged cDNA fragments in the scRNA-seq library. These factors could lead to high dropout rates of CD56-aligned reads from the final gene×cell tallies in the PBMC 3K expression matrix [94,103].

**Figure 4.**
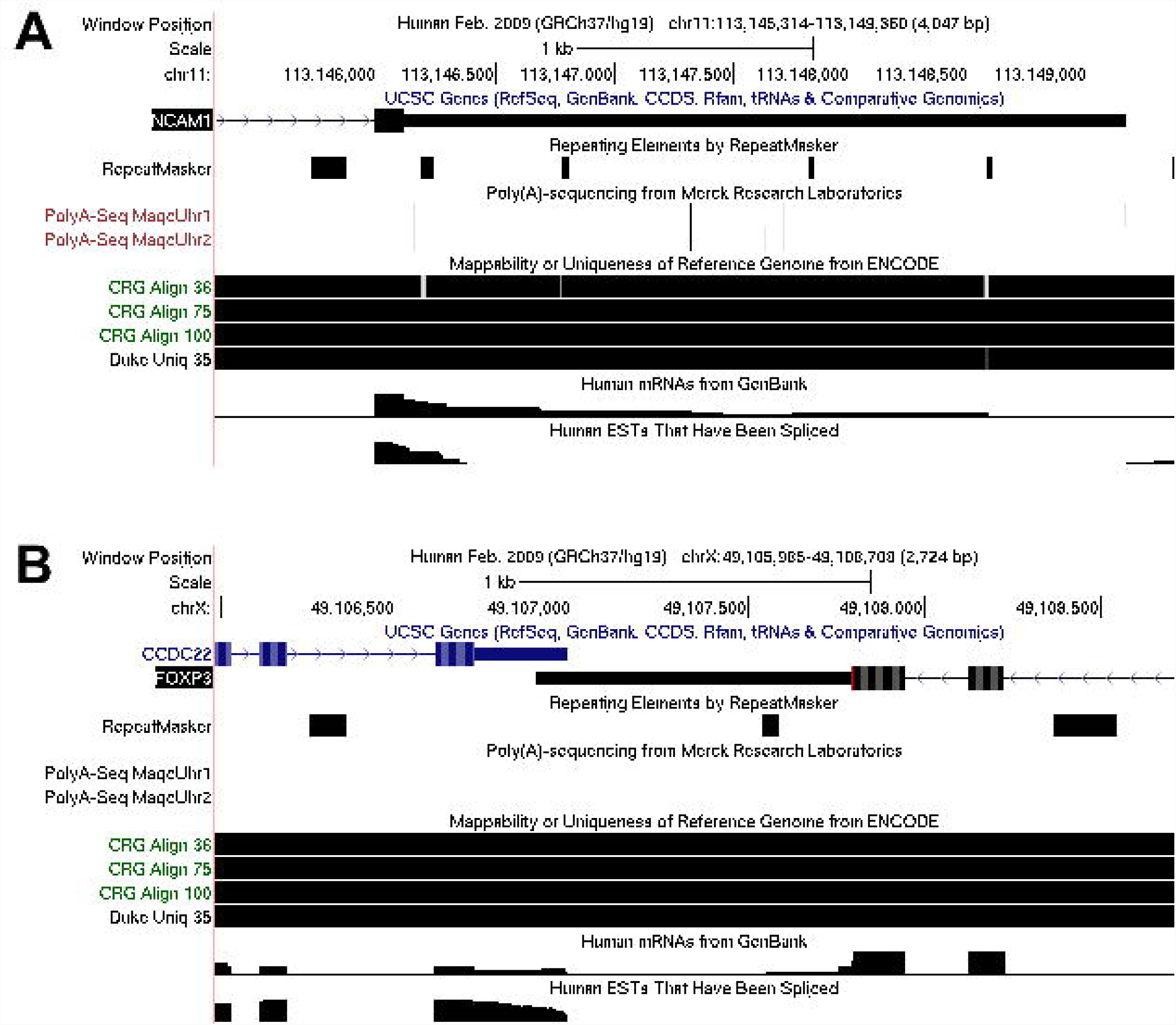
Genomic features at the 3’UTR locus of CD56(NCAM1) and FOXP3 in hg19. Genomic 3-Kb windows in the UCSC Genome Browser spanning across the canonical 3’ UTRs of human genes (A) CD56(NCAM1) and (B) FOXP3. Genomic tracks depict, from top to bottom: intron/exon encoding regions based on UCSC annotations, repetitive elements recognized by RepeatMasker, inferred poly-A^+^ insertion sites based on 3’ end alignments of poly-A^+^ RNA-seq data using MAQC Universal Human Reference samples, mappability scores of sliding window *k*-mers based on CRG Alignability (36-, 75-, and 100-nt) and Duke Uniqueness (100-nt) pipelines, density tracks for overlapping mRNAs in Genbank, and density tracks for overlapping spliced expressed sequence tags (EST) in Genbank.

Another example of the mapping challenges in scRNA-seq can be seen in the case of FOXP3. Previously known as scurfin, FOXP3 is a critical transcription factor that commits CD4+ T cells towards immune tolerance functions (known as Tregs) and, to similar extent, CD8+ T cells towards immunosuppressive roles (known as Tsups) [56,59,65,67,70]. The 3’ UTR of FOXP3 in the human reference genome is ∼900 bp long and contains one (CTGGGG)_n_ single-repeat element. The repeat is located roughly 650 bp upstream from the only poly-A^+^ insertion site annotated for FOXP3. It lies within 130 bp from the 3’ end of the mRNA which extends the longest into the 3’ UTR (390 bp) among the Genbank-annotated FOXP3 transcripts. However, the single-repeat element within the 3’ UTR of FOXP3, unlike those in the 3’ UTR of CD56, can be aligned uniquely from sequenced k-mers as small as 36 bp [94,103]. This feature effectively rules out the possibility that low counts for FOXP3 in the PBMC 3K dataset (11 individual cells, 1 UMI each) are due to high dropouts of reads from its 3’ UTR that are not uniquely mappable.

Instead, we propose that the low detection rate of FOXP3 transcripts may stem from the ongoing regulation of FOXP3 mRNA stability in leukocytes. The 3’ UTR of FOXP3 is targeted by SMHD1, a constitutively expressed ribonucleolytic enzyme endogenous to immune cells *in vivo*, in particular resting T cells [104,105]. Critically, these findings suggest that recognizing subsets of CD4+ Tregs or CD8+ Tsups in peripheral blood would be extremely difficult based only on FOXP3 read counts from scRNA-seq data. Instead, predicting FOXP3-positive cells would be best achieved by surveying scRNA-seq data for gene transcripts highly correlated with antibody-based detection of FOXP3 protein in circulating cells, such as CD25(IL2R) or CD127(IL7R) [56,59,65,67,68,70]. In the PBMC 3K data, which is derived from peripheral blood from a healthy subject, this alternative signature matches to small pockets within the CD4+ and CD8+ populations in cell majors A and C, respectively (Fig. 3E and S1).

## Discussion

In developing the SALSA workflow, we sought to define a systematic approach that enhances replicability and reproducibility in the analysis of scRNA-seq data. Currently, most mainstream approaches resolve cell cluster from single cell data by testing the whole contents of the gene-cell matrix; SALSA demonstrates that it is more advantageous to prioritize facultative genes for inferential testing. SALSA introduces parametric stratification of detected genes in the gene-cell matrix to determine which genes most likely exhibit variable expression among individual cells. Here we tackle four core challenges key to devising a general purpose, LSA-based scRNA-seq workflow which arise from discrepancies in how different research groups handle and generate their scRNA-seq data. Firstly, we leverage a portioning approach to assess if sequenced reads carrying the same cell barcode are representatives of a single-cell transcriptome. We then constructed a framework to define if mRNA transcripts mapped to certain genes are representative of variably or endogenously expressed ones within an individual biological specimen. Subsequently, these values are used to infer how single-cell transcriptomes converge into classes of overarching cell profiles which translate into DEG-driven transcriptional signatures. Ultimately, SALSA deduces minimal sets of reproducible and cell profile-specific DEGs, or Profiler genes, suited for biomarker-based experimental validation.

Implementation of SALSA repurposes the structure of single-cell transcriptomics data into a latent variable extraction problem. In return, this approach confers two major advantages to scRNA-seq data analysis: it reduces the number of genes needed for inferential testing and increases statistical robustness. By handling the data in this way, we introduce expression transformants that lend single-cell DEG extraction with statistical compliance. In addition, with this approach we can utilise highly efficient and widely available multivariate analyses algorithms that rely on linearity and homoscedasticity assumptions, such as ANOVAs and hierarchical clustering. SALSA bypasses the more common step of gene×cell expression matrix assembly where empty gene×cell blocks are artificially filled in with zeros [34,37,43,52]. Considering that these empty blocks amount to well over 90% of all possible gene×cell crossings in any previously reported scRNA-seq benchmark datasets [15,36], doing without aggressively zero-inflated gene×cell expression matrices generates smaller gene expression files. When combined with highly efficient SVD-driven algorithms for sparse matrix imputation and latent variable extraction, the SALSA workflow cuts down on the computational footprint and memory allocation typically required for scRNA-seq data post-processing. Critically, t [27,44,52,92,106]. hese statistical enhancements provide biomarker outputs that are testable on the bench because SALSA prioritises genes that can be verified using conventional validation approaches.

SALSA, like other recently reported tools for unguided bioinformatic inferencing of hierarchical associations in biology [107,108], leverages the concepts behind latent semantic analysis (LSA) methods: the concepts of “genes” and “single cells” found in scRNA-seq data can be thought of as interchangeable with the concepts of “terms” and “documents” in natural language processing algorithms [109-112]. Both SALSA and LSA perform eigenvalue optimization driven by explicit count data in ultra-sparse matrices using “local” measures of relative frequency for gene/term counts in a cell/document, such as UMI-per-thousand (UPT) or count-per-document total scaling. Additionally, both SALSA and LSA implement “global” weight systems to adjust for the overall frequency of a gene/term vs. all order gene/terms detected anywhere. SALSA and LSA differ in how they compile the preponderance of detected gene/term counts into a useful statistical kernel. In LSA, “global” weights are used to adjust for the incidence of terms throughout a *known* corpus of documents ahead of inferential testing, with the inverse document frequency being the most commonly used “global” weighting statistic. In the SALSA workflow, UPT values from a sequencing round are estimated from counts of UMIs which must initially be deduplicated, disambiguated, and ascribed to an *unknown* number of single cells. These values are inferred from a pool of observed barcodes and then must be empirically transformed into linearized and normally distributed metrics via generalized linear modelling [9,45,113-120]. Critically, such metrics are uniquely pertinent to each experimental dataset being analysed. It follows that by retaining statistical idiosyncrasy for separate scRNA-seq runs, SALSA imposes a replicability constraint: if a facultative gene is to score as a “true” DEG, it must do so across multiple biological specimens despite batch effects. Additionally, a “true” DEG must re-score as a consensus DEG free of replication bias once data from any within-replicate DEGs are re-tested as an all-at-once ensemble. This concept will be explored in more detail in future publications.

We show that using SALSA, we can isolate the portions harbouring the most informative subset of DEGs in an expression matrix. Conversely, we show that non-facultative genes are detrimental to DEG extraction from both an experimental and bioinformatics perspective. On one side, retaining low-coverage genes risks admitting UMI data from contaminant or anecdotal templates, such as poly-A^+^ tracks from gDNA debris or ambient RNA in cell suspensions. On the other side, including ubiquitously transcribed RNA reads in differential expression analysis, such as reads aligned to rRNA or housekeeping gene mRNAs, biases statistical significance testing for genes with higher total UMI counts but small log-fold expression differences. By introducing matrix focusing into scRNA-seq data processing, the SALSA workflow prioritizes data within the dynamic range of expression measurements. By doing so based on parametric distribution fitting, statistical inferences on focused expression matrices yield results that can be replicated systematically across groups using equivalent datasets. In sum, the SALSA workflow reduces the dimensionality of gene-cell matrices and curtails oversampling of measurement errors, which boosts statistical power of differential expression analyses.

Unlike other scRNA-seq bioinformatic approaches based on barcode filtering using spline functions, SALSA can make reproducible calls by using parametric distribution functions as a criterion to decide if the number of sequenced reads carrying the same barcode are representative of a single-cell transcriptome relative to the total from all other detected barcodes within the same library. Additionally, SALSA can make reproducible calls on whether transcripts in single-cell transcriptomes, mapped to specific genes, are representative of variably expressed genes or endogenously expressed genes within individual biological specimens. The SALSA workflow helps us infer which single-cell transcriptomes fall into classes of overarching cell profiles; each of these profiles will exhibit a transcriptional signature driven by a core group of DEGs. Using SALSA, we unveiled distinct cell profiles represented at comparable rates, and driven by expressed gene modules unique to each cell; we termed these Profiler genes. The sets of Profiler genes represented top candidates for biomarker-based experimental validation. To allow for success on the bench, genes categorised as Profilers must have expression that is mutually exclusive among multiple cell subpopulations. Ontological analysis can then be performed on the robust set of DEGs identified by SALSA to extract pathway-level insights about identified cell profiles based on replicate scRNA-seq assays.

In our view, PBMC 3K represents an important resource and reference, because it harbors one of the densest UMI data counts among publicly available scRNA-seq datasets. However, this dataset also highlights pervasive limitations in the biochemistry of scRNA-seq library assembly despite its relatively rich UMI coverage rate per cell. In tackling the PBMC 3K dataset with SALSA, we were surprised to find that the classic markers CD56 and FOXP3 were unreliable transcriptional markers to discriminate peripheral blood cell types in scRNA-seq data because of their poor representation in the expression matrix. Our analysis suggests CD56 exhibited a high bioinformatic dropout rate and the absence of FOXP3 message could be explained by quick turn over by endogenous ribonucleases. Critically, if it is the intrinsic features of the 3’ UTR sequences of CD56 and FOXP3 transcripts that are to blame for their poor detection rates, our findings would imply that increasing the depth of sequencing will not enrich CD56 or FOXP3 read alignments meaningfully. Quite opposite, our findings suggest that generating additional coverage chasing for more CD56 or FOXP3 events would instead enrich for gDNA debris or ambient RNA reads, diluting the differences between true expression signal and background contaminants across the board.

In all, we put forth the SALSA workflow, a parsimonious approach to scRNA-seq data mining, to demonstrate how reoccurring and experimentally testable sets of cell type-specific DEGs can be compiled across scRNA-seq datasets from independent biological specimens. In that respect, being able to recognize phenotypic expression patterns by stratifying detected genes, at a fraction of the computational fingerprint of other pipelines, represents the largest asset of the SALSA workflow. We believe data focusing strategies like those used in SALSA will finally allow data integration from replicative experimental designs to become mainstream in single-cell ‘omics – while fostering research reproducibility in the field along the way [121]. Regarding resource allocation, SALSA extracts the maximum amount of bench-verifiable information from data-sparse scRNA-seq datasets by recognizing latent relations among statistically robust genes, which evoke cell type-specific functions, and does so without relying on lineage-specific transcripts being known in advance. These advances in scRNA-seq data mining open the door to making multi-platform confirmatory testing and orthogonal hardware comparisons financially feasible. Future work will demonstrate how to use SALSA to integrate single-cell transcriptomics datasets across independent specimens and scRNA-seq platforms, especially on how to deduce batch-insensitive sets of reproducible DEGs that belong to underlying cell types. Bottom line: SALSA was developed to elevate biological insight and maximize data profits from expensive single cell technologies, hopefully encouraging their dissemination in a world with limiting resources.

## Online Methods

### Publicly available PBMC 3K dataset from 10X Genomics

Count-level scRNA-seq data for peripheral mononuclear blood cells of a healthy human subject retrieved from a commercially available frozen stock [62] is available for download from 10X Genomics (https://support.10xgenomics.com/single-cell-gene-expression/datasets). Data was generated by sequencing a scRNA-seq library produced with 10X GemCode technology and run in a single Illumina NextSeq 500 high-output flow cell (∼69,000 raw reads per deplexed barcode). Further details on scRNA-seq library assembly process, sequencing data acquisition, and single-cell barcode discrimination pipelines are available in the original publication by Zheng et al. [36]. For our analyses, we used a curated version of the PBMC 3K dataset as consensus, with UMI alignments to ∼17K annotated transcripts among 2,700 single-cell barcodes inferred by the 10X Genomics’ CellRanger 1.1.0 pipeline; the consensus curated version of the PBMC 3K dataset is available online for download, courtesy of Rahul Satija’s research group, at: https://s3-us-west-2.amazonaws.com/10x.files/samples/cell/pbmc3k/pbmc3k_filtered_gene_bc_matrices.tar.gz.

### Parametric focusing of gene-cell expression matrices

A diagram of matrix focusing process is depicted in Fig. 1B-D. Briefly, we identified “best-guess” single-cell barcodes and facultative genes within each specimen based on the empirical distribution function of total UMIs each by regression analysis to a parametric 2-component Weibull-Frechét mixture model and heavy-tailed Frechét model to project an upper bound of inlier total UMI coverage. Further details on the statistical treatment of total UMI count data for matrix focusing are available in the Supplementary Methods section.

### Implementation of Single-cell Amalgamation by Latent Semantic Analysis (SALSA)

A diagram of the SALSA statistical workflow is depicted in Fig. 1E. Briefly, expression levels of individual genes (Ensembl annotation) in individual cells were calculated as the normalized rate of deduplicated and uniquely aligned UMIs per thousand total UMIs per cell. Similarly, bulk reference expression levels for individual genes (Ensembl annotation) were calculated as the normalized UPT rate for all UMIs in the library combined. Bulk UPT rates of genes were used to extract a best-fit parametric threshold distribution of expression intensities from the exponential family, then used in combination with single-cell UPT rates to determine a linear predictor of expression intensity in normalized space ***B***(***θ*)** via generalized linear modelling of bulk UPT rates with a distribution-matched natural parameter and a corresponding canonical link function [119]. Next, we defined prospective cell clusters via IRLBA-driven clustering [44] of mean linear predictor estimates «***B***(**θ**) », and carried out prospective differential expression analysis between them via the LSTNR method [45] with adjustments for representation rates of gene transcripts per cluster along the workflow (as depicted in Fig. 1E). Gene resolution weights of genes corresponded to the cumulative hazard of gene-wise significance scores from two-way ANOVA testing of the GLM linear predictor scores. Within-cluster gene representation rates are defined as the ratio of cells with aligned UMIs vs. total cells within a cluster for each gene. Statistical tests of differential gene expression were performed using double-weighed two-way ANOVA model (gene×cluster blocks) of log2-transformed fold changes in single-cell UPT rates (Log_2_FC) relative to gene-wise bulk UPT rates. We refer to double-weighed ANOVA tests as those combining the product of gene resolution weights and within-cluster gene representation rates into a single weight parameter for two-way ANOVA testing of Log_2_FC values. Gene-wise significance of Log2FC variation based on double-weighed ANOVA tests were adjusted by the Benjamini-Hochberg method for multiple comparisons [122]. All metrics and statistical analyses were carried out using JMP® 13.0.0 64-bit statistical software (SAS, Cary, NC).

### Stratification of significantly expressed genes

Significant genes (SGs) were identified as those with significantly double-weighted ANOVA differences (FDR adj. p<0.05). As a reproducibility benchmark, we refer to LSTNRs as the subset of SGs in which a minimum practical effect size δ_Log2FC_>(δ_Effect_= 0.3×σ_SSR_) was detected and at least one prospective cell cluster exhibits average Log_2_FC signal greater than a library-wide minimum measurement precision defined as the 95% Tolerance Interval of gene×cluster residuals among SGs (i.e. with SNR>1). Afterwards, cell assignments into groups are refined by IRLBA re-clustering of into cell majors based on mean linear predictor estimates "***B***(**θ**)" from LSTNR genes only. Differentially expressed genes (DEGs) are the subset of LSTNRs with significant differences (FDR adj. p<0.05) in Log_2_FC values among cell majors by single-weighted (representation only, disregarding resolution) ANOVA tests and a cardinal number (≥*N-*1 for *N* cell majors) of *post hoc* significant Log_*2*_FC pairwise differences (Δ_Log2FC_) between clusters (Student’s t-test p<0.05). We deem differentially expressed genes with reproducible expectation estimates (DEGREEs) as the subset of DEGs for which the number of Δ_Log2FC_ > 95% TI_SSR_ is also cardinal. Finally, the subset of DEGREEs that exhibit significant differences in Log_2_FC values among clusters based on unweighted ANOVA tests (i.e. disregarding representation and resolution), with δ_Log2FC_>(δ_Effect_= 0.3×σ_SSR_), SNR>1, and cardinal number of *post hoc* significant Δ_Log2FC_ > 95% TI_SSR_ between cell majors, corresponds to Profiler genes.

## Supporting information

Supplementary Discussion and Methods

## Author Contributions

OAL conceptualized and implemented the analytical methodology described in the study, performed data analysis, and conceived data visualization strategies; OAL and KSM wrote the initial manuscript drafts leading to submission; and BP compiled and curated scRNA-seq data. JL and HY supervised the selection of scRNA-seq datasets to validate and test the statistical approach described herein. All authors contributed in the design of the study and interpretation of the results, as well as revised, read and approved the submitted version.

## Acknowledgments

We would like to thank Jay Shendure and Junyue Cao (University of Washington, USA) and members of the Yao Lab (NIEHS, NIH, USA), Woychik Lab (NIEHS, NIH, USA), and Oliver Lab (NIDDK, NIH, USA) for helpful discussions around the concepts in this paper. We also thank Pierre Bushel at the Biostatistics and Computational Biology Group (NIEHS, NIH, USA) and Suzanne Matos in the Bell Lab (NIEHS, NIH, USA) for their critical review of this manuscript ahead of submission. We thank Peter Koopman (University of Queensland, Australia) for his helpful advice and guidance throughout this project.

## Funding

This research was supported by the Intramural Research Program (ES102965 to HHCY) of the NIH, National Institute of Environmental Health Sciences.

## Conflict of Interest

The authors declare that the research was conducted in the absence of any commercial or financial relationships that could be construed as a potential conflict of interest.

## Supplementary Figures

Figure S1. Schematic of total UMI coverage bound estimates for single-cell barcode regimes using P_C_-P_D_ mixture model parameters.

Figure S2. Topographs for 55 landmark and supplemental expression markers of blood cell types, as detected in the PBMC 3K dataset.

Figure S3. Profiler genes enriched in cell majors B, F and G of the PBMC 3K dataset, based on multinomial logistic regression of weighed expression rates.

## Supplementary Tables

**Table S1. List of 166 genes in the PBMC 3K dataset with multi-count UMI alignments (4 or more counts) among gene×cell data-positive expression matrix fields.**

## References

1. Mortazavi A, Williams BA, McCue K, Schaeffer L, Wold B. Mapping and quantifying mammalian transcriptomes by RNA-Seq. Nat Methods. 2008;5(7):621–628. doi:10.1038/nmeth.1226. PubMed PMID: 18516045.

2. Oshlack A, Robinson MD, Young MD. From RNA-seq reads to differential expression results. Genome Biol. 2010;11(12):220. doi:10.1186/gb-2010-11-12-220. PubMed PMID: 21176179; PubMed Central PMCID: PMC3046478.

3. Roy NC, Altermann E, Park ZA, McNabb WC. A comparison of analog and Next-Generation transcriptomic tools for mammalian studies. Brief Funct Genomics. 2011;10(3):135–150. doi:10.1093/bfgp/elr005. PubMed PMID: 21389008.

4. Huynh NPT, Zhang B, Guilak F. High-depth transcriptomic profiling reveals the temporal gene signature of human mesenchymal stem cells during chondrogenesis. FASEB J. 2018:fj201800534R. doi:10.1096/fj.201800534R. PubMed PMID: 29985644.

5. Li B, Dorrell C, Canaday PS, Pelz C, Haft A, Finegold M, et al. Adult Mouse Liver Contains Two Distinct Populations of Cholangiocytes. Stem Cell Reports. 2017;9(2):478–489. doi:10.1016/j.stemcr.2017.06.003. PubMed PMID: 28689996; PubMed Central PMCID: PMCPMC5549808.

6. Oikawa T, Wauthier E, Dinh TA, Selitsky SR, Reyna-Neyra A, Carpino G, et al. Model of fibrolamellar hepatocellular carcinomas reveals striking enrichment in cancer stem cells. Nat Commun. 2015;6:8070. doi:10.1038/ncomms9070. PubMed PMID: 26437858; PubMed Central PMCID: PMCPMC4600730.

7. Cloonan N, Forrest AR, Kolle G, Gardiner BB, Faulkner GJ, Brown MK, et al. Stem cell transcriptome profiling via massive-scale mRNA sequencing. Nat Methods. 2008;5(7):613–619. doi:10.1038/nmeth.1223. PubMed PMID: 18516046.

8. Gong B, Wang C, Su Z, Hong H, Thierry-Mieg J, Thierry-Mieg D, et al. Transcriptomic profiling of rat liver samples in a comprehensive study design by RNA-Seq. Sci Data. 2014;1:140021. doi:10.1038/sdata.2014.21. PubMed PMID: 25977778; PubMed Central PMCID: PMCPMC4322565.

9. Li J, Bushel PR. EPIG-Seq: extracting patterns and identifying co-expressed genes from RNA-Seq data. BMC Genomics. 2016;17:255. doi:10.1186/s12864-016-2584-7. PubMed PMID: 27004791; PubMed Central PMCID: PMC4804494.

10. McClelland KS, Bell K, Larney C, Harley VR, Sinclair AH, Oshlack A, et al. Purification and Transcriptomic Analysis of Mouse Fetal Leydig Cells Reveals Candidate Genes for Specification of Gonadal Steroidogenic Cells. Biol Reprod. 2015;92(6):145. doi:10.1095/biolreprod.115.128918. PubMed PMID: 25855264.

11. van den Brink SC, Sage F, Vertesy A, Spanjaard B, Peterson-Maduro J, Baron CS, et al. Single-cell sequencing reveals dissociation-induced gene expression in tissue subpopulations. Nat Methods. 2017;14(10):935–936. doi:10.1038/nmeth.4437. PubMed PMID: 28960196.

12. Cao J, Cusanovich DA, Ramani V, Aghamirzaie D, Pliner HA, Hill AJ, et al. Joint profiling of chromatin accessibility and gene expression in thousands of single cells. Science. 2018. doi:10.1126/science.aau0730. PubMed PMID: 30166440.

13. Cao J, Packer JS, Ramani V, Cusanovich DA, Huynh C, Daza R, et al. Comprehensive single-cell transcriptional profiling of a multicellular organism. Science. 2017;357(6352):661–667. doi:10.1126/science.aam8940. PubMed PMID: 28818938; PubMed Central PMCID: PMCPMC5894354.

14. Klein AM, Mazutis L, Akartuna I, Tallapragada N, Veres A, Li V, et al. Droplet barcoding for single-cell transcriptomics applied to embryonic stem cells. Cell. 2015;161(5):1187–1201. doi:10.1016/j.cell.2015.04.044. PubMed PMID: 26000487; PubMed Central PMCID: PMCPMC4441768.

15. Macosko EZ, Basu A, Satija R, Nemesh J, Shekhar K, Goldman M, et al. Highly Parallel Genome-wide Expression Profiling of Individual Cells Using Nanoliter Droplets. Cell. 2015;161(5):1202–1214. doi:10.1016/j.cell.2015.05.002. PubMed PMID: 26000488; PubMed Central PMCID: PMCPMC4481139.

16. Picelli S, Bjorklund AK, Faridani OR, Sagasser S, Winberg G, Sandberg R. Smart-seq2 for sensitive full-length transcriptome profiling in single cells. Nat Methods. 2013;10(11):1096–1098. doi:10.1038/nmeth.2639. PubMed PMID: 24056875.

17. Picelli S, Faridani OR, Bjorklund AK, Winberg G, Sagasser S, Sandberg R. Full-length RNA-seq from single cells using Smart-seq2. Nat Protoc. 2014;9(1):171–181. doi:10.1038/nprot.2014.006. PubMed PMID: 24385147.

18. Rosenberg AB, Roco CM, Muscat RA, Kuchina A, Sample P, Yao Z, et al. Single-cell profiling of the developing mouse brain and spinal cord with split-pool barcoding. Science. 2018;360(6385):176–182. doi:10.1126/science.aam8999. PubMed PMID: 29545511.

19. Trapnell C, Cacchiarelli D, Grimsby J, Pokharel P, Li S, Morse M, et al. The dynamics and regulators of cell fate decisions are revealed by pseudotemporal ordering of single cells. Nat Biotechnol. 2014;32(4):381–386. doi:10.1038/nbt.2859. PubMed PMID: 24658644; PubMed Central PMCID: PMCPMC4122333.

20. Satija R, Farrell JA, Gennert D, Schier AF, Regev A. Spatial reconstruction of single-cell gene expression data. Nat Biotechnol. 2015;33(5):495–502. doi:10.1038/nbt.3192. PubMed PMID: 25867923; PubMed Central PMCID: PMCPMC4430369.

21. Briggs JA, Weinreb C, Wagner DE, Megason S, Peshkin L, Kirschner MW, et al. The dynamics of gene expression in vertebrate embryogenesis at single-cell resolution. Science. 2018;360(6392). doi:10.1126/science.aar5780. PubMed PMID: 29700227; PubMed Central PMCID: PMCPMC6038144.

22. Farrell JA, Wang Y, Riesenfeld SJ, Shekhar K, Regev A, Schier AF. Single-cell reconstruction of developmental trajectories during zebrafish embryogenesis. Science. 2018;360(6392). doi:10.1126/science.aar3131. PubMed PMID: 29700225.

23. Alles J, Karaiskos N, Praktiknjo SD, Grosswendt S, Wahle P, Ruffault PL, et al. Cell fixation and preservation for droplet-based single-cell transcriptomics. BMC Biol. 2017;15(1):44. doi:10.1186/s12915-017-0383-5. PubMed PMID: 28526029; PubMed Central PMCID: PMCPMC5438562.

24. Wu YE, Pan L, Zuo Y, Li X, Hong W. Detecting Activated Cell Populations Using Single-Cell RNA-Seq. Neuron. 2017;96(2):313–329 e316. doi:10.1016/j.neuron.2017.09.026. PubMed PMID: 29024657.

25. Attar M, Sharma E, Li S, Bryer C, Cubitt L, Broxholme J, et al. A practical solution for preserving single cells for RNA sequencing. Sci Rep. 2018;8(1):2151. doi:10.1038/s41598-018-20372-7. PubMed PMID: 29391536; PubMed Central PMCID: PMCPMC5794922.

26. Butler A, Hoffman P, Smibert P, Papalexi E, Satija R. Integrating single-cell transcriptomic data across different conditions, technologies, and species. Nat Biotechnol. 2018;36(5):411–420. doi:10.1038/nbt.4096. PubMed PMID: 29608179.

27. Haghverdi L, Lun ATL, Morgan MD, Marioni JC. Batch effects in single-cell RNA-sequencing data are corrected by matching mutual nearest neighbors. Nat Biotechnol. 2018;36(5):421–427. doi:10.1038/nbt.4091. PubMed PMID: 29608177; PubMed Central PMCID: PMCPMC6152897.

28. Hashimshony T, Wagner F, Sher N, Yanai I. CEL-Seq: single-cell RNA-Seq by multiplexed linear amplification. Cell Rep. 2012;2(3):666–673. doi:10.1016/j.celrep.2012.08.003. PubMed PMID: 22939981.

29. Islam S, Kjallquist U, Moliner A, Zajac P, Fan JB, Lonnerberg P, et al. Characterization of the single-cell transcriptional landscape by highly multiplex RNA-seq. Genome Res. 2011;21(7):1160–1167. doi:10.1101/gr.110882.110. PubMed PMID: 21543516; PubMed Central PMCID: PMCPMC3129258.

30. Jaitin DA, Kenigsberg E, Keren-Shaul H, Elefant N, Paul F, Zaretsky I, et al. Massively parallel single-cell RNA-seq for marker-free decomposition of tissues into cell types. Science. 2014;343(6172):776–779. doi:10.1126/science.1247651. PubMed PMID: 24531970; PubMed Central PMCID: PMCPMC4412462.

31. Marcy Y, Ishoey T, Lasken RS, Stockwell TB, Walenz BP, Halpern AL, et al. Nanoliter reactors improve multiple displacement amplification of genomes from single cells. PLoS Genet. 2007;3(9):1702–1708. doi:10.1371/journal.pgen.0030155. PubMed PMID: 17892324; PubMed Central PMCID: PMCPMC1988849.

32. Ramskold D, Luo S, Wang YC, Li R, Deng Q, Faridani OR, et al. Full-length mRNA-Seq from single-cell levels of RNA and individual circulating tumor cells. Nat Biotechnol. 2012;30(8):777–782. doi:10.1038/nbt.2282. PubMed PMID: 22820318; PubMed Central PMCID: PMCPMC3467340.

33. Sasagawa Y, Nikaido I, Hayashi T, Danno H, Uno KD, Imai T, et al. Quartz-Seq: a highly reproducible and sensitive single-cell RNA sequencing method, reveals non-genetic gene-expression heterogeneity. Genome Biol. 2013;14(4):R31. doi:10.1186/gb-2013-14-4-r31. PubMed PMID: 23594475; PubMed Central PMCID: PMCPMC4054835.

34. Shalek AK, Satija R, Adiconis X, Gertner RS, Gaublomme JT, Raychowdhury R, et al. Single-cell transcriptomics reveals bimodality in expression and splicing in immune cells. Nature. 2013;498(7453):236–240. doi:10.1038/nature12172. PubMed PMID: 23685454; PubMed Central PMCID: PMCPMC3683364.

35. Tang F, Barbacioru C, Wang Y, Nordman E, Lee C, Xu N, et al. mRNA-Seq whole-transcriptome analysis of a single cell. Nat Methods. 2009;6(5):377–382. doi:10.1038/nmeth.1315. PubMed PMID: 19349980.

36. Zheng GX, Terry JM, Belgrader P, Ryvkin P, Bent ZW, Wilson R, et al. Massively parallel digital transcriptional profiling of single cells. Nat Commun. 2017;8:14049. doi:10.1038/ncomms14049. PubMed PMID: 28091601; PubMed Central PMCID: PMCPMC5241818 L.M., D.A.M., S.Y.N., M.S.L., P.W.W., C.M.H., R.B., A.W., K.D.N., T.S.M. and B.J.H. are employees of 10x Genomics.

37. Zilionis R, Nainys J, Veres A, Savova V, Zemmour D, Klein AM, et al. Single-cell barcoding and sequencing using droplet microfluidics. Nat Protoc. 2017;12(1):44–73. doi:10.1038/nprot.2016.154. PubMed PMID: 27929523.

38. Dong YY, Shen S. Testing for Rank Invariance or Similarity in Program Evaluation. Review of Economics and Statistics. 2018;100(1):78–85. doi:10.1162/REST_a_00686. PubMed PMID: WOS:000426598900007.

39. Frandsen BR, Lefgren LJ. Testing Rank Similarity. Review of Economics and Statistics. 2018;100(1):86–91. doi:10.1162/REST_a_00675. PubMed PMID: WOS:000426598900008.

40. Stanley HE, Amaral LAN, Gopikrishnan P, Ivanov PC, Keitt TH, Plerou V. Scale invariance and universality: organizing principles in complex systems. Physica A. 2000;281(1-4):60–68. doi:Doi 10.1016/S0378-4371(00)00195-3. PubMed PMID: WOS:000087741700007.

41. Nair J, Wierman A, Zwart B. Tail-robust scheduling via limited processor sharing. Performance Evaluation. 2010;67(11):978–995. doi:10.1016/j.peva.2010.08.012. PubMed PMID: WOS:000283808600002.

42. Cont R. Empirical properties of asset returns: stylized facts and statistical issues. Quantitative Finance. 2001;1(2):223–236. doi:10.1080/713665670. PubMed PMID: WOS:000208488800006.

43. Wu AR, Neff NF, Kalisky T, Dalerba P, Treutlein B, Rothenberg ME, et al. Quantitative assessment of single-cell RNA-sequencing methods. Nat Methods. 2014;11(1):41–46. doi:10.1038/nmeth.2694. PubMed PMID: 24141493; PubMed Central PMCID: PMCPMC4022966.

44. Baglama J, Reichel L. Augmented implicitly restarted Lanczos bidiagonalization methods. Siam Journal on Scientific Computing. 2005;27(1):19–42. doi:10.1137/04060593x. PubMed PMID: WOS:000232354900002.

45. Lozoya OA, Santos JH, Woychik RP. A Leveraged Signal-to-Noise Ratio (LSTNR) Method to Extract Differentially Expressed Genes and Multivariate Patterns of Expression From Noisy and Low-Replication RNAseq Data. Front Genet. 2018;9:176. doi:10.3389/fgene.2018.00176. PubMed PMID: 29868123; PubMed Central PMCID: PMCPMC5964166.

46. Raj A, Peskin CS, Tranchina D, Vargas DY, Tyagi S. Stochastic mRNA synthesis in mammalian cells. PLoS Biol. 2006;4(10):e309. doi:10.1371/journal.pbio.0040309. PubMed PMID: 17048983; PubMed Central PMCID: PMCPMC1563489.

47. Larsson AJM, Johnsson P, Hagemann-Jensen M, Hartmanis L, Faridani OR, Reinius B, et al. Genomic encoding of transcriptional burst kinetics. Nature. 2019;565(7738):251–254. doi:10.1038/s41586-018-0836-1. PubMed PMID: 30602787.

48. Hausser J, Mayo A, Keren L, Alon U. Central dogma rates and the trade-off between precision and economy in gene expression. Nat Commun. 2019;10(1):68. doi:10.1038/s41467-018-07391-8. PubMed PMID: 30622246.

49. Liu Y, Beyer A, Aebersold R. On the Dependency of Cellular Protein Levels on mRNA Abundance. Cell. 2016;165(3):535–550. doi:10.1016/j.cell.2016.03.014. PubMed PMID: 27104977.

50. Moulana A, Scanteianu A, Jones D, Stern AD, Bouhaddou M, Birtwistle M. Gene-Specific Predictability of Protein Levels from mRNA Data in Humans. bioRxiv. 2018:399816. doi:10.1101/399816.

51. Mohammadi S, Davila-Velderrain J, Kellis M, Grama A. DECODE-ing sparsity patterns in single-cell RNA-seq. bioRxiv. 2018:241646. doi:10.1101/241646.

52. Zhang MJ, Ntranos V, Tse D. One read per cell per gene is optimal for single-cell RNA-Seq. bioRxiv. 2018:389296. doi:10.1101/389296.

53. Chauhan AK. Human CD4(+) T-Cells: A Role for Low-Affinity Fc Receptors. Front Immunol. 2016;7:215. doi:10.3389/fimmu.2016.00215. PubMed PMID: 27313579; PubMed Central PMCID: PMCPMC4887501.

54. Gustafson MP, Lin Y, Maas ML, Van Keulen VP, Johnston PB, Peikert T, et al. A method for identification and analysis of non-overlapping myeloid immunophenotypes in humans. PloS one. 2015;10(3):e0121546. doi:10.1371/journal.pone.0121546. PubMed PMID: 25799053; PubMed Central PMCID: PMCPMC4370675.

55. Henson SM, Riddell NE, Akbar AN. Properties of end-stage human T cells defined by CD45RA re-expression. Curr Opin Immunol. 2012;24(4):476–481. doi:10.1016/j.coi.2012.04.001. PubMed PMID: 22554789.

56. Hu Z, Jujjavarapu C, Hughey JJ, Andorf S, Lee HC, Gherardini PF, et al. MetaCyto: A Tool for Automated Meta-analysis of Mass and Flow Cytometry Data. Cell Rep. 2018;24(5):1377–1388. doi:10.1016/j.celrep.2018.07.003. PubMed PMID: 30067990.

57. Mahnke YD, Brodie TM, Sallusto F, Roederer M, Lugli E. The who’s who of T-cell differentiation: human memory T-cell subsets. Eur J Immunol. 2013;43(11):2797–2809. doi:10.1002/eji.201343751. PubMed PMID: 24258910.

58. Sallusto F, Lenig D, Forster R, Lipp M, Lanzavecchia A. Two subsets of memory T lymphocytes with distinct homing potentials and effector functions. Nature. 1999;401(6754):708–712. doi:10.1038/44385. PubMed PMID: 10537110.

59. Zhao C, Davies JD. A peripheral CD4+ T cell precursor for naive, memory, and regulatory T cells. J Exp Med. 2010;207(13):2883–2894. doi:10.1084/jem.20100598. PubMed PMID: 21149551; PubMed Central PMCID: PMCPMC3005223.

60. Tan C, Taylor AA, Coburn MZ, Marino JH, Van De Wiele CJ, Teague TK. Ten-color flow cytometry reveals distinct patterns of expression of CD124 and CD126 by developing thymocytes. BMC Immunol. 2011;12:36. doi:10.1186/1471-2172-12-36. PubMed PMID: 21689450; PubMed Central PMCID: PMCPMC3130696.

61. Zola H, Flego L, Weedon H. Expression of IL-4 receptor on human T and B lymphocytes. Cell Immunol. 1993;150(1):149–158. doi:10.1006/cimm.1993.1186. PubMed PMID: 7688267.

62. AllCells®. 2019 [01/25/2019]. Available from: https://www.allcells.com/products/peripheral-blood-mononuclear-cells-pbmc.

63. Lin A, Lore K. Granulocytes: New Members of the Antigen-Presenting Cell Family. Front Immunol. 2017;8:1781. doi:10.3389/fimmu.2017.01781. PubMed PMID: 29321780; PubMed Central PMCID: PMCPMC5732227.

64. Pyzik M, Rath T, Lencer WI, Baker K, Blumberg RS. FcRn: The Architect Behind the Immune and Nonimmune Functions of IgG and Albumin. Journal of Immunology. 2015;194(10):4595–4603. doi:10.4049/jimmunol.1403014. PubMed PMID: 25934922; PubMed Central PMCID: PMCPMC4451002.

65. Churlaud G, Pitoiset F, Jebbawi F, Lorenzon R, Bellier B, Rosenzwajg M, et al. Human and Mouse CD8(+)CD25(+)FOXP3(+) Regulatory T Cells at Steady State and during Interleukin-2 Therapy. Front Immunol. 2015;6:171. doi:10.3389/fimmu.2015.00171. PubMed PMID: 25926835; PubMed Central PMCID: PMCPMC4397865.

66. Du X, Tang Y, Xu H, Lit L, Walker W, Ashwood P, et al. Genomic profiles for human peripheral blood T cells, B cells, natural killer cells, monocytes, and polymorphonuclear cells: comparisons to ischemic stroke, migraine, and Tourette syndrome. Genomics. 2006;87(6):693–703. doi:10.1016/j.ygeno.2006.02.003. PubMed PMID: 16546348.

67. Khattri R, Cox T, Yasayko SA, Ramsdell F. An essential role for Scurfin in CD4+CD25+ T regulatory cells. Nat Immunol. 2003;4(4):337–342. doi:10.1038/ni909. PubMed PMID: 12612581.

68. Liu W, Putnam AL, Xu-Yu Z, Szot GL, Lee MR, Zhu S, et al. CD127 expression inversely correlates with FoxP3 and suppressive function of human CD4+ T reg cells. J Exp Med. 2006;203(7):1701–1711. doi:10.1084/jem.20060772. PubMed PMID: 16818678; PubMed Central PMCID: PMCPMC2118339.

69. Ahlers JD, Belyakov IM. Memories that last forever: strategies for optimizing vaccine T-cell memory. Blood. 2010;115(9):1678–1689. doi:10.1182/blood-2009-06-227546. PubMed PMID: 19903895; PubMed Central PMCID: PMCPMC2920202.

70. Yagi H, Nomura T, Nakamura K, Yamazaki S, Kitawaki T, Hori S, et al. Crucial role of FOXP3 in the development and function of human CD25+CD4+ regulatory T cells. Int Immunol. 2004;16(11):1643–1656. doi:10.1093/intimm/dxh165. PubMed PMID: 15466453.

71. Karnell JL, Dimasi N, Karnell FG, 3rd, Fleming R, Kuta E, Wilson M, et al. CD19 and CD32b differentially regulate human B cell responsiveness. Journal of Immunology. 2014;192(4):1480–1490. doi:10.4049/jimmunol.1301361. PubMed PMID: 24442430; PubMed Central PMCID: PMCPMC3918864.

72. Bednar F, Song C, Bardi G, Cornwell W, Rogers TJ. Cross-desensitization of CCR1, but not CCR2, following activation of the formyl peptide receptor FPR1. Journal of Immunology. 2014;192(11):5305–5313. doi:10.4049/jimmunol.1302983. PubMed PMID: 24778447; PubMed Central PMCID: PMCPMC4035699.

73. Hambleton J, Weinstein SL, Lem L, DeFranco AL. Activation of c-Jun N-terminal kinase in bacterial lipopolysaccharide-stimulated macrophages. Proc Natl Acad Sci U S A. 1996;93(7):2774–2778. PubMed PMID: 8610116; PubMed Central PMCID: PMCPMC39708.

74. Shi GX, Harrison K, Han SB, Moratz C, Kehrl JH. Toll-like receptor signaling alters the expression of regulator of G protein signaling proteins in dendritic cells: implications for G protein-coupled receptor signaling. Journal of Immunology. 2004;172(9):5175–5184. PubMed PMID: 15100254.

75. Wakabayashi Y, Kobayashi M, Akashi-Takamura S, Tanimura N, Konno K, Takahashi K, et al. A protein associated with toll-like receptor 4 (PRAT4A) regulates cell surface expression of TLR4. Journal of Immunology. 2006;177(3):1772–1779. PubMed PMID: 16849487.

76. Mandl M, Schmitz S, Weber C, Hristov M. Characterization of the CD14++CD16+ monocyte population in human bone marrow. PloS one. 2014;9(11):e112140. doi:10.1371/journal.pone.0112140. PubMed PMID: 25369328; PubMed Central PMCID: PMCPMC4219836.

77. Sanyal R, Polyak MJ, Zuccolo J, Puri M, Deng L, Roberts L, et al. MS4A4A: a novel cell surface marker for M2 macrophages and plasma cells. Immunol Cell Biol. 2017;95(7):611–619. doi:10.1038/icb.2017.18. PubMed PMID: 28303902.

78. Kowal K, Osada J, Zukowski S, Dabrowska M, Dubuske L, Bodzenta-Lukaszyk A. Expression of interleukin 4 receptors in bronchial asthma patients who underwent specific immunotherapy. Ann Allergy Asthma Immunol. 2004;93(1):68–75. doi:10.1016/S1081-1206(10)61449-4. PubMed PMID: 15281474.

79. Baig A, Bao X, Wolf M, Haslam RJ. The platelet protein kinase C substrate pleckstrin binds directly to SDPR protein. Platelets. 2009;20(7):446–457. doi:10.3109/09537100903137314. PubMed PMID: 19852682.

80. Brinkmann V, Zychlinsky A. Neutrophil extracellular traps: is immunity the second function of chromatin? J Cell Biol. 2012;198(5):773–783. doi:10.1083/jcb.201203170. PubMed PMID: 22945932; PubMed Central PMCID: PMCPMC3432757.

81. Cullen PJ. Calcium signalling: the ups and downs of protein kinase C. Curr Biol. 2003;13(18):R699–701. PubMed PMID: 13678606.

82. Diaz-Godinez C, Carrero JC. The state of art of neutrophil extracellular traps in protozoan and helminthic infections. Biosci Rep. 2019;39(1). doi:10.1042/BSR20180916. PubMed PMID: 30498092; PubMed Central PMCID: PMCPMC6328873.

83. Lian L, Wang Y, Flick M, Choi J, Scott EW, Degen J, et al. Loss of pleckstrin defines a novel pathway for PKC-mediated exocytosis. Blood. 2009;113(15):3577–3584. doi:10.1182/blood-2008-09-178913. PubMed PMID: 19190246; PubMed Central PMCID: PMCPMC2668855.

84. Liverani E, Mondrinos MJ, Sun S, Kunapuli SP, Kilpatrick LE. Role of Protein Kinase C-delta in regulating platelet activation and platelet-leukocyte interaction during sepsis. PloS one. 2018;13(4):e0195379. doi:10.1371/journal.pone.0195379. PubMed PMID: 29617417; PubMed Central PMCID: PMCPMC5884571.

85. Walz A, Dewald B, von Tscharner V, Baggiolini M. Effects of the neutrophil-activating peptide NAP-2, platelet basic protein, connective tissue-activating peptide III and platelet factor 4 on human neutrophils. J Exp Med. 1989;170(5):1745–1750. PubMed PMID: 2681518; PubMed Central PMCID: PMCPMC2189513.

86. Gustafsson K, Ingelsten M, Bergqvist L, Nystrom J, Andersson B, Karlsson-Parra A. Recruitment and activation of natural killer cells in vitro by a human dendritic cell vaccine. Cancer Res. 2008;68(14):5965–5971. doi:10.1158/0008-5472.CAN-07-6494. PubMed PMID: 18632652.

87. Martinet J, Dufeu-Duchesne T, Bruder Costa J, Larrat S, Marlu A, Leroy V, et al. Altered functions of plasmacytoid dendritic cells and reduced cytolytic activity of natural killer cells in patients with chronic HBV infection. Gastroenterology. 2012;143(6):1586–1596 e1588. doi:10.1053/j.gastro.2012.08.046. PubMed PMID: 22960656.

88. Pokkali S, Das SD, Selvaraj A. Differential upregulation of chemokine receptors on CD56 NK cells and their transmigration to the site of infection in tuberculous pleurisy. FEMS Immunol Med Microbiol. 2009;55(3):352–360. doi:10.1111/j.1574-695X.2008.00520.x. PubMed PMID: 19159432.

89. Hanna J, Bechtel P, Zhai YF, Youssef F, McLachlan K, Mandelboim O. Novel insights on human NK cells’ immunological modalities revealed by gene expression profiling. Journal of Immunology. 2004;173(11):6547–6563. doi:DOI 10.4049/jimmunol.173.11.6547. PubMed PMID: WOS:000225307500009.

90. Poli A, Michel T, Theresine M, Andres E, Hentges F, Zimmer J. CD56bright natural killer (NK) cells: an important NK cell subset. Immunology. 2009;126(4):458–465. doi:10.1111/j.1365-2567.2008.03027.x. PubMed PMID: 19278419; PubMed Central PMCID: PMCPMC2673358.

91. Romee R, Foley B, Lenvik T, Wang Y, Zhang B, Ankarlo D, et al. NK cell CD16 surface expression and function is regulated by a disintegrin and metalloprotease-17 (ADAM17). Blood. 2013;121(18):3599–3608. doi:10.1182/blood-2012-04-425397. PubMed PMID: 23487023; PubMed Central PMCID: PMCPMC3643761.

92. Picelli S. Single-cell RNA-sequencing: The future of genome biology is now. Rna Biology. 2017;14(5):637–650. doi:10.1080/15476286.2016.1201618. PubMed PMID: WOS:000402095400017.

93. Coffey AJ, Kokocinski F, Calafato MS, Scott CE, Palta P, Drury E, et al. The GENCODE exome: sequencing the complete human exome. Eur J Hum Genet. 2011;19(7):827–831. doi:10.1038/ejhg.2011.28. PubMed PMID: 21364695; PubMed Central PMCID: PMCPMC3137498.

94. Derrien T, Estelle J, Marco Sola S, Knowles DG, Raineri E, Guigo R, et al. Fast computation and applications of genome mappability. PloS one. 2012;7(1):e30377. doi:10.1371/journal.pone.0030377. PubMed PMID: 22276185; PubMed Central PMCID: PMCPMC3261895.

95. Chung BY, Hardcastle TJ, Jones JD, Irigoyen N, Firth AE, Baulcombe DC, et al. The use of duplex-specific nuclease in ribosome profiling and a user-friendly software package for Ribo-seq data analysis. RNA. 2015;21(10):1731–1745. doi:10.1261/rna.052548.115. PubMed PMID: 26286745; PubMed Central PMCID: PMCPMC4574750.

96. O’Neil D, Glowatz H, Schlumpberger M. Ribosomal RNA depletion for efficient use of RNA-seq capacity. Curr Protoc Mol Biol. 2013; Chapter 4:Unit 4 19. doi:10.1002/0471142727.mb0419s103. PubMed PMID: 23821444.

97. Yoshihama M, Uechi T, Asakawa S, Kawasaki K, Kato S, Higa S, et al. The human ribosomal protein genes: sequencing and comparative analysis of 73 genes. Genome Res. 2002;12(3):379–390. doi:10.1101/gr.214202. PubMed PMID: 11875025; PubMed Central PMCID: PMCPMC155282.

98. Zhao W, He X, Hoadley KA, Parker JS, Hayes DN, Perou CM. Comparison of RNA-Seq by poly (A) capture, ribosomal RNA depletion, and DNA microarray for expression profiling. BMC Genomics. 2014;15:419. doi:10.1186/1471-2164-15-419. PubMed PMID: 24888378; PubMed Central PMCID: PMCPMC4070569.

99. Zhulidov PA, Bogdanova EA, Shcheglov AS, Vagner LL, Khaspekov GL, Kozhemyako VB, et al. Simple cDNA normalization using kamchatka crab duplex-specific nuclease. Nucleic Acids Res. 2004;32(3):e37. doi:10.1093/nar/gnh031. PubMed PMID: 14973331; PubMed Central PMCID: PMCPMC373426.

100. Mayr C. Regulation by 3’-Untranslated Regions. Annu Rev Genet. 2017;51:171–194. doi:10.1146/annurev-genet-120116-024704. PubMed PMID: 28853924.

101. Matoulkova E, Michalova E, Vojtesek B, Hrstka R. The role of the 3’ untranslated region in post-transcriptional regulation of protein expression in mammalian cells. Rna Biology. 2012;9(5):563–576. doi:10.4161/rna.20231. PubMed PMID: 22614827.

102. Cheng J, Maier KC, Avsec Z, Rus P, Gagneur J. Cis-regulatory elements explain most of the mRNA stability variation across genes in yeast. RNA. 2017;23(11):1648–1659. doi:10.1261/rna.062224.117. PubMed PMID: 28802259; PubMed Central PMCID: PMCPMC5648033.

103. Kent WJ, Sugnet CW, Furey TS, Roskin KM, Pringle TH, Zahler AM, et al. The human genome browser at UCSC. Genome Res. 2002;12(6):996–1006. doi:10.1101/gr.229102. Article published online before print in May 2002. PubMed PMID: 12045153; PubMed Central PMCID: PMC186604.

104. Kim YC, Kim KK, Yoon J, Scott DW, Shevach EM. SAMHD1 Posttranscriptionally Controls the Expression of Foxp3 and Helios in Human T Regulatory Cells. Journal of Immunology. 2018;201(6):1671–1680. doi:10.4049/jimmunol.1800613. PubMed PMID: 30104243; PubMed Central PMCID: PMCPMC6125212.

105. Ruffin N, Brezar V, Ayinde D, Lefebvre C, Schulze Zur Wiesch J, van Lunzen J, et al. Low SAMHD1 expression following T-cell activation and proliferation renders CD4+ T cells susceptible to HIV-1. AIDS. 2015;29(5):519–530. doi:10.1097/QAD.0000000000000594. PubMed PMID: 25715102; PubMed Central PMCID: PMCPMC4342413.

106. Qiu X, Rahimzamani A, Wang L, Mao Q, Durham T, McFaline-Figueroa JL, et al. Towards inferring causal gene regulatory networks from single cell expression measurements. bioRxiv. 2018:426981. doi:10.1101/426981.

107. Cong Y, Chan YB, Ragan MA. A novel alignment-free method for detection of lateral genetic transfer based on TF-IDF. Sci Rep. 2016;6:30308. doi:10.1038/srep30308. PubMed PMID: 27453035; PubMed Central PMCID: PMCPMC4958984.

108. Moussa M, Mandoiu, II. Single cell RNA-seq data clustering using TF-IDF based methods. BMC Genomics. 2018;19(Suppl 6):569. doi:10.1186/s12864-018-4922-4. PubMed PMID: 30367575.

109. Luhn HP. The Automatic Creation of Literature Abstracts. Ibm Journal of Research and Development. 1958;2(2):159–165. doi:DOI 10.1147/rd.22.0159. PubMed PMID: WOS:A1958WY42200007.

110. Salton G, Buckley C. Term-Weighting Approaches in Automatic Text Retrieval. Information Processing & Management. 1988;24(5):513–523. doi:Doi 10.1016/0306-4573(88)90021-0. PubMed PMID: WOS:A1988Q414000001.

111. Sparck-Jones K. A statistical interpretation of term specificity and its application in retrieval. Journal of Documentation. 2004;60(5):493–502. doi:10.1108/00220410410560573. PubMed PMID: WOS:000224871900002.

112. Wu HC, Luk RWP, Wong KF, Kwok KL. Interpreting TF-IDF term weights as making relevance decisions. Acm Transactions on Information Systems. 2008;26(3). doi:Artn 13 10.1145/1361684.1361686. PubMed PMID: WOS:000257028500002.

113. Aitkin M, Clayton D. The Fitting of Exponential, Weibull and Extreme Value Distributions to Complex Censored Survival Data Using GLIM. Journal of the Royal Statistical Society Series C (Applied Statistics). 1980;29(2):156–163. doi:10.2307/2986301.

114. Bullard JH, Purdom E, Hansen KD, Dudoit S. Evaluation of statistical methods for normalization and differential expression in mRNA-Seq experiments. BMC Bioinformatics. 2010;11:94. doi:10.1186/1471-2105-11-94. PubMed PMID: 20167110; PubMed Central PMCID: PMC2838869.

115. Finotello F, Di Camillo B. Measuring differential gene expression with RNA-seq: challenges and strategies for data analysis. Brief Funct Genomics. 2015;14(2):130–142. doi:10.1093/bfgp/elu035. PubMed PMID: 25240000.

116. Hansen KD, Wu Z, Irizarry RA, Leek JT. Sequencing technology does not eliminate biological variability. Nat Biotechnol. 2011;29(7):572–573. doi:10.1038/nbt.1910. PubMed PMID: 21747377; PubMed Central PMCID: PMC3137276.

117. Law CW, Chen Y, Shi W, Smyth GK. voom: Precision weights unlock linear model analysis tools for RNA-seq read counts. Genome Biol. 2014;15(2):R29. doi:10.1186/gb-2014-15-2-r29. PubMed PMID: 24485249; PubMed Central PMCID: PMC4053721.

118. Li J, Tibshirani R. Finding consistent patterns: a nonparametric approach for identifying differential expression in RNA-Seq data. Stat Methods Med Res. 2013;22(5):519–536. doi:10.1177/0962280211428386. PubMed PMID: 22127579; PubMed Central PMCID: PMC4605138.

119. Nelder JA, Wedderburn RW. Generalized Linear Models. Journal of the Royal Statistical Society Series a-General. 1972;135(3):370-+. doi:Doi 10.2307/2344614. PubMed PMID: WOS:A1972N770800027.

120. Oberg AL, Bot BM, Grill DE, Poland GA, Therneau TM. Technical and biological variance structure in mRNA-Seq data: life in the real world. BMC Genomics. 2012;13:304. doi:10.1186/1471-2164-13-304. PubMed PMID: 22769017; PubMed Central PMCID: PMC3505161.

121. Ioannidis JP. Why most published research findings are false. PLoS Med. 2005;2(8):e124. doi:10.1371/journal.pmed.0020124. PubMed PMID: 16060722; PubMed Central PMCID: PMC1182327.

122. Benjamini Y, Hochberg Y. Controlling the False Discovery Rate: A Practical and Powerful Approach to Multiple Testing. Journal of the Royal Statistical Society Series B (Methodological). 1995;57(1):289–300.

