## Supplementary Discussion and Methods for "Patterns, Profiles, and Parsimony: dissecting transcriptional signatures from minimal single-cell RNA-seq output with SALSA"

#### Experimental errors and distributional features for single-cell RNAseq data with different barcoding strategies

A challenge in calculating read counts in all modalities of RNAseq arises when any two reads that map to the same genomic coordinates are detected in the same library. One solution is to append a unique sequence “handle” to reverse transcribed copies of mRNA molecules during cDNA assembly; such a unique molecular identifier (UMI) is usually added during oligonucleotide synthesis within reverse transcription primers by subsequent rounds of single random base pair addition. Introducing UMIs in reverse transcription primers ensures the same UMI is common to all PCR copies from a single cDNA template; later, UMIs help distinguish whether two reads with highly homologous sequences that share the same genome alignment were: a) derived from two distinct cDNA molecules; or b) degenerate PCR copies from the same cDNA template that differ due to base pair integration errors.

In single-cell RNAseq, reads are pooled by their presumed cell of origin (which is designated by barcode sequences) and mapped uniquely to the reference genome. Then, reads are tallied per mapped gene and assigned to individual fields in a gene×cell matrix describing the total read counts mapped to each individual gene (rows) under each captured cell barcode (columns), which we refer to as the “hash matrix” denoted by ***R***. For that reason, it is critical that the tally in ***R*** is representative of the initial pool of unique mRNA molecules, and that “double-counting” is prevented – which can occur when cDNA templates giving rise to matching PCR copies cannot be traced back to their original mRNA molecule. It follows that when the number of starting mRNA molecules is low, the probability increases that many fragments in the final sequencing library will be PCR copies derived from a single cDNA template. Also, if the sequencing library consists of randomly cleaved UMI-encoding fragments of cDNA copies amassed by PCR, then sequenced reads may be detected that map to slightly different genomic ranges (i.e. PCR copies of each initial cDNA template cleaved into slightly different fragments) while sharing the same UMI (i.e. derived from the same mRNA molecule).

In this context, use of UMIs is particularly relevant to the assembly of single-cell RNAseq libraries, which originate from limiting amounts of mRNA molecules per cell – with only a small fraction of those contributing to cDNA synthesis reactions. The limited yield of cDNA templates derives in low-complexity libraries with a limited mass of distinct fragments. Given mass and size constraints that dsDNA libraries must meet to load successfully onto sequencing flow cells, low-complexity libraries usually undergo successive rounds of purification steps for size selection (e.g. SPRI-based size selection) and removal of oligonucleotide adapter sequences and/or PCR primers. Doing so, however, carries with it an inevitable loss of library mass; the usual remedy against those losses is adding PCR amplification rounds in between purification steps. The hope is to increase the probability of having each initial mRNA template represented in the final sequencing library more than once – by creating multiple copies of fragments derived from the original cDNA templates – to counterbalance mass losses during library purification. For those reasons, the usual outcome after sequencing a low-complexity single-cell RNAseq library is a pool of dsDNA reads composed almost exclusively (>95%) of PCR copies from a small set of UMI-bearing cDNA templates – representing only fragments from cDNA copies that were retained across the entire specimen-to-sequencer hands-on workflow.

As in any other NGS technique, DNA artifacts may contaminate a sample and, on occasion, integrate UMI handles or NGS sequencing adapters needed for sequencing at any step in the specimen-to-sequencer hands-on workflow. In the case of RNAseq, these artifacts may include primer dimers, ambient RNA, or impurities in PCR consumables; specifically, for single-cell RNAseq, mRNA molecules or nucleic acid debris from broken cells that disperse in suspension media are not only templates for cDNA synthesis but also artifacts, because their appended barcodes cannot trace back to one cell of origin. If (when) they occur, any barcoded DNA artifacts become sequencing-ready dsDNA templates in the library that may add to the total counts of detected reads.

In bulk RNAseq assays, where a single barcode annotates transcripts from thousands of cells at once, contributions by DNA artifacts are usually outweighed by the overall diversity of distinct mRNA molecules present. However, the detriment of such artifacts to single-cell RNAseq data is much more substantial. Unfortunately, most bioinformatic pipelines to populate ***R*** matrices from UMI-deplexed reads do not discriminate between UMIs shared by many sequenced fragments and UMIs detected only once – in other words, UMIs are counted all the same in data post-processing, regardless of whether they were added to the library at the start (e.g. mRNA reverse transcription) or in between steps (e.g. dsDNA contaminants) of the specimen-to-sequencer hands-on workflow. In effect, this also means that UMI-harboring artifacts can impact each field in ***R*** matrices in at least two ways: by distributing heterogeneously across barcodes, or by harboring barcodes of their own. These matters are complicated further by the fact that total UMI counts accrued from individual cell barcodes across detected genes (or vice versa) vary, meaning that an equal number of dsDNA counts added to different cell barcodes does not have the same impact on their expression measurement uncertainty when their total counts are different – i.e. once normalized, the “statistical error” from dsDNA artifacts is different. For all those reasons, a pre-filtering step is needed *even before quantification of single-cell gene expression* to recognize *bona fide* barcodes and UMIs representative of mRNA molecules tractable to their individual cells of origin, while discarding UMI-harboring artifacts that map to the reference genome successfully – some of which do so by mere chance.

In principle, one may anticipate whether the apportionment of dsDNA artifacts in data from a single-cell RNAseq library will lean towards an all-around “smeared UMI noise” mixed with *bona fide* UMIs across multiple cell barcodes, or towards dsDNA artifacts harboring barcodes of their own. Such bias would depend on both the experimental approach to single-cell indexing and the quality of the single cell suspension used for library assembly; presumably, this inference may also work the other way around: the distribution of UMIs per detected barcode in a single-cell RNAseq data set is also a probabilistic fingerprint of the experimental technique and sample conditions used to synthesize a sequenced single-cell RNAseq library. This *a priori* inference is key to devising a frequentist pre-filtering approach to minimize the impact of dsDNA artifact counts to single-cell expression analysis. To do so, recognizing how barcoding noise and artifacts manifest in overall UMI distributions is key, and depends on which one of (at least) two broadly classified methods is chosen to couple single cells to individual barcodes: by droplet-based encapsulation or combinatorial indexing.

Currently, the most affordable droplet-based encapsulation systems rely on dual split-flow microfluidic devices that feed two separate incoming aqueous suspensions – i.e. cells and barcoding oligonucleotides – into a junction and down a single channel; the coalescing flow is met downstream by a third incoming cross-flow of an immiscible fluid – e.g. oil – that “pinches” the aqueous stream to form single micelles of equal volume. To assign a shared barcode to all transcripts captured from the same cell, oligonucleotides with the same underlying barcode must be packaged into single delivery depots and appended to cDNA copies of RNA templates during reverse transcription. Such barcode depots are built by synthesizing reverse transcription primers onto immobilizing substrates, or carriers, that include randomly integrated nucleotide sequences using successive single-base addition rounds. In this manner, the number of nucleotides in barcodes determines the number of distinct sequences that can be generated; hence, the probability of two or more carrier particles sharing the same random barcode lessens as more nucleotides are stringed onto fewer immobilizing carriers (since the number of possible nucleotide sequences increases) and all but vanishes once the number of random barcodes is greater than the number of supplied carrier particles. Afterwards, random UMI sequences are added to all carriers at once, followed by functional reverse transcription primers – e.g. poly(dT) sequences complementary to poly-A tails found in mRNA transcripts – such that all oligonucleotide molecules immobilized to each single carrier share the same barcode, yet each contains a UMI of its own, and can prime cDNA synthesis from RNA templates. In this manner, each cDNA copy that enters the library assembly workflow is tagged with information about the carrier that captured its source RNA molecule but remains distinguishable from cDNA copies of RNA molecules that hybridized to other oligonucleotides on the same carrier. Today, the most widely used carriers consist of surface-functionalized porous resins or oligo-laden hydrogels and are suspended in buffered solutions containing reverse transcription enzymes and detergents to lyse the cells that carriers are to be encapsulated with. The purpose is to ensure that once lysed within droplets, single cells release RNA templates to be used for compartmentalized cDNA synthesis reactions on the surface of co-encapsulated barcode carriers, and that the necessary ingredients to carry out reverse transcription are available inside every droplet.

When using a dual split-flow microfluidic device to generate single-cell RNAseq libraries, the probability of initiating uniquely barcoded cDNA synthesis reactions using RNA templates from a single cell is bounded by the probability that a flowing single and unique barcode carrier coincides with the contents of a single cell upon encapsulation – which requires that cells and carriers both reach the junction separately, one of each at a time, and whole. Assuming the arrivals of non-lysed single cells and barcode carriers to an encapsulation junction are both mutually independent Poisson-distributed events, the maximum theoretical probability of singlet collision events (i.e. the probability of a single cell coinciding with a single carrier inside the same droplet) is equal to $1/e^{2}$, or no more than ∼13.5% of all generated droplets containing at least one of the two, and occurs when the average feed rates of cells and carriers match. Conversely, the theoretical minimum proportion of droplets with either no carriers or only carriers, no cells or only cells, multiple carriers that coincide with more than one cell in the same droplet (“multiplets”) or vice versa, or empty (i.e. only aqueous media from cell and carrier suspensions) is at least ~86.5% of all droplets produced – i.e. over 6 times the maximum number of expectable singlets. In practical terms, the product of a single-cell encapsulation run is a heterogeneous colloid that consists of mostly multiplets (whose barcodes tag the contents of many cells each) and “empty” droplets (whose contents can only supply trace DNA contaminants, ambient RNA, or cell debris to cDNA synthesis reactions) along with a much smaller fraction of true singlets overall; therefore, most barcodes detected upon sequencing are to be discarded during data processing. Even then, it is relatively simple to recognize barcoded artifacts in a droplet-based single-cell RNAseq library – whether they randomly map to a reference genome (with the lowest UMI total counts overall) or originate as cDNA copies of ambient RNA from lysed cells (with diluted UMI total counts) – as they amount to most of the detected barcodes, and have the lowest sequencing representation; therefore, barcoded artifacts fall at the lower end of quantile plots for total UMI counts per barcode. Conversely, because their proportion in the library is much smaller but have larger and relatively stable total UMI counts each, “true” single-cell barcodes show up as a “late” plateau in quantile plots. Multiplets are found further up the barcode representation range and exhibit a progression in total UMI counts away from the single-cell plateau in the quantile plot.

Often, supply rates of cells and carriers (or their concentrations) are manipulated to minimize production of multiplets and obtain a final colloid composed of only singlets or empty droplets, such that each sequenced barcode can be assigned to either class during post-processing simply based on having many or few total UMIs, respectively. However, using mismatched supply rates of cells and carriers lowers singlet collision rates dramatically, thereby increasing the proportion of empty droplet barcodes that harbor low UMI totals each (as they tag only DNA artifacts or ambient RNA). This approach can become a double-edged sword during data processing: in the one hand, the distribution of UMI totals per barcode from empty droplets can be leveraged as a minimum of read counts expected per singlet; in the other hand, an overrepresentation of empty droplets in the colloid risks leading to a library composed almost exclusively of empty barcodes, and producing a dataset of only artifacts. This trade-off can only be evaluated empirically, as transcription rates in single cells vary between experimental replicates, cell types, and biological specimens. Also, the ability to distinguish between singlet and empty barcodes based on their total UMI counts depends on how many artifacts like ambient RNA molecules and cell debris are present in the mixing flows. Therefore, the purity of buffers and reagents, handling of biological specimens, and integrity of single-cell suspensions is critical to discriminating between empty barcodes and singlets among an unknown set and number of supplied barcode carriers.

Barcoding in combinatorial indexing (or “split-pooling”) starts with *in situ* cDNA synthesis reactions, such that cells or nuclei are used as encapsulation units in the library assembly workflow. In principle, this method relies on reverse transcription primers and enzymes having molecular weights smaller than size exclusion limits of organellar membranes, so they can diffuse in and reach RNA templates; conversely, RNA templates larger than those exclusion limits cannot exit. Supplying reagents and nucleotides in excess can also impose an osmotic gradient, thereby enhancing this unidirectional exchange. When performed on isolated nuclei or fixed cells, this transport mechanism is passive, thus requiring no microfluidics devices. Once inside, reagents and primers can be activated to integrate index sequences within cDNA molecules inside individual cells; furthermore, if the integrity of cells or organelles is maintained, multiple rounds of indexing can be carried out where indexed cells from one multi-well plate can be retrieved, resuspended, and delivered at random into different multi-well plates for additional indexing rounds. This iterative process results in compounded barcodes, consisting of combinations of known indices appended in successive split-pool rounds, that cDNA templates from a single cell all share and can be decoded during sequenced data post-processing. Therefore, the number of possible index combinations determines a finite number of single-cell compounded barcodes that can turn up in a single-cell sequencing library and depends exclusively on the number of split-pool rounds carried out while using a set number of indices. In practical terms, this also means that the number of split-pool rounds can be chosen in advance to generate more index combinations than the number of cells to be sequenced, all but guaranteeing that the probability of multiple cells sharing the same compounded barcode is negligible.

Above all, the main advantage of combinatorial indexing techniques is that the underlying indices and the number of compounded barcodes they can assemble are fixed and known in advance. This feature of combinatorial indexing also suggests that if the numbers of single cells to be indexed is not only less, but of a similar order of magnitude than the number of barcodes as well, then artifacts are unlikely to harbor barcodes of their own. Instead, in combinatorial indexing, UMIs appended to artifacts will most likely smear across all barcodes as a background signal that contributes to total UMI counts across the board. In that sense, artifacts behave as a detection threshold to distinguish barcodes whose total UMI make-up is predominantly derived from single-cell RNA templates instead of trace contaminants, PCR concatemers, or ambient RNA from lysed cells. Such a threshold can be estimated by parametric fits of total UMI counts per barcode to a truncated probability function and would be applicable to all barcodes at once. For that reason, any barcodes with sequencing representation over the threshold qualify as single cells, comprise most of the detected barcodes, and appear as an upshot in quantile plots of total UMI counts per barcode (since sequencing representation is usually similar among true single cells vs. artifacts).

Still, the desired number of sequenced cells is a critical parameter to consider when devising an experimental design, whether a droplet-based or a combinatorial indexing approach is chosen. As the number of PCR cycles required to generate sufficient library mass for sequencing necessarily rises with lower cell numbers, PCR fidelity errors accumulate more in specimens containing fewer cells; thus, the base integration errors in the library increase, and degeneracies in barcode and UMI sequence replication start piling up – hindering the purpose of UMI technology altogether. In other words, the probability of misrepresenting UMI sequences is directly linked to the accumulated number of PCR cycles needed to assemble and sequence the library itself. In practice, PCR error “catch-up” in low-complexity single-cell RNAseq libraries means that deeper sequencing in hopes of raising the total UMI counts per barcode may be inherently risky – or even detrimental – to a faithful interpretation of differential gene expression between single cells within a sequenced specimen. To what extent does oversaturation of library complexity in a sequencing run shifts towards production of reads carrying spurious barcodes and UMIs that mismatch those of their original templates (and resemble “new” data) is still a problem for NGS technologies across the board. In lieu of robust corrective measures to interpret “true” UMIs apart from degenerate ones *a posteriori*, it is our view that larger cell numbers, use of ERCC spike-ins, or empirical determination of *quantum sufficit* sequencing depths for different types of specimens all remain the most practical strategies to optimizing bioinformatic analytical pipelines and enhancing reproducible differential expression analysis at the single-cell level.

### Supplementary Methods

#### Unifying probabilistic mixture model for single-cell sequencing depth parameterization

Understanding how artifact representation differs between droplet- v. combinatorial-based techniques has important implications to inferring whether UMI-containing reads represent cells or artifacts. As discussed previously, the expectable noise fingerprints in UMI representation of artifacts are almost opposite between droplet-based and combinatorial-based techniques from a statistical perspective: whereas artifacts tend to accrue their own barcodes in droplet-based libraries, they contribute background reads to all barcodes indiscriminately in combinatorial indexing libraries instead. Hence, artifact representation differences between techniques should *in principle* be clearly discernible by displaying quantile plots of total UMI counts per barcode.

In practice, though, capturing such stark contrasts between separate libraries assembled with either droplet- or combinatorial-based techniques is experimentally possible only if the biological specimen to be sequenced is truly dissociated into single cells, which do not clump together over time, while ensuring that ambient RNA from premature cell lysis in suspension medium is minimal. This is rarely the case, and often results in quantile plots of total UMI counts per barcode that resemble a mix of both the droplet-based and split-pool-based methods, with a bias towards the actual technique used. In that sense, total UMI distributions in single-cell RNAseq libraries can be thought of as mixtures of two extreme cases: the theoretical droplet-based and combinatorial-based profiles. Therefore, a 2-distribution parametric mixture model is the simplest statistical model to fit empirical distributions of total UMIs from single-cell RNAseq libraries experimentally produced using either technique.

Consider a single-cell RNAseq library ***R*** described by a *B*×*G* tensor of sequenced UMIs, each encoding one of *B* unique barcodes and uniquely mapped to one of *G* total genes represented in a sequenced library. Thus, ***R*** is the total set of unique sequencing read counts in the library, or “hash matrix”, such that

$$\boldsymbol{R=}R_{ij}$$

describes (in Einstein notation) a tally of nonduplicate templates detected in the library under the *i*^th^ captured barcode and mapped to the *j*^th^ gene in the reference genome. Thus, given an all-ones column vector $\mathcal{X}_{k}=1$ of size *G*×1,

$$b_{i}=R_{ik}\mathcal{X}_{k}$$

represents the total counted and uniquely identified sequences in the library carrying the *i*^th^ barcode – i.e. the total UMI counts in the *i*^th^ barcode.

Now, let us define a probabilistic mixture model of $b_{i}$ with $N\geq n$parametric probability functions, such that

$$w_{n}\text{δ}_{nk}=1$$

and

$$\text{P}\left( b_{i}\leq x \right)=w_{n}\text{P}_{n}\text{δ}_{nk}+\varepsilon_{n}\text{δ}_{nk}$$

given the set of positive weights $w_{n}$ for individual parametric probability functions $\text{P}_{n}$, where $\text{δ}_{nk}$ is the Kronecker delta, and $\varepsilon_{n}$ are regression error terms. Therefore, a 2-distribution parametric mixture model of theoretical droplet-based combinatorial-based probability functions is given by

$$\text{P}\left( b_{i}\leq x \right)=w_{\text{D}}\cdot\text{P}_{\text{D}}+w_{\text{C}}\cdot\text{P}_{\text{C}}+\varepsilon_{D}{+\varepsilon}_{C}$$

where

$$w_{\text{D}}=1-w_{\text{C}}$$

and $\text{P}_{\text{D}}$ and $\text{P}_{\text{C}}$ denote probability functions for theoretical (i.e. without multiplets) droplet-based and combinatorial-based libraries, respectively.

If we assume that the theoretical droplet-based library consists mostly of barcodes with low UMI counts assigned to artifacts that transitions afterwards into a small proportion of barcodes with higher UMI counts for singlets, then its theoretical distribution $\text{P}_{\text{D}}$ for total UMIs per barcode $b_{i}$ can be represented by a Weibull probability function bounded at $x>0$ such that

$$\text{P}_{\text{D}}\left( b_{i}\leq x \right) \sim\text{Wb}\left( x;\beta,\alpha\right)=\frac{\beta}{\alpha}\cdot\left( \frac{x}{\alpha} \right)^{\beta-1}\cdot exp\left[ -\left( \frac{x}{\alpha} \right)^{\beta} \right]$$

where $\text{P}_{\text{D}}$ refers to a theoretical droplet-based library, and $\beta>0$ and $\alpha>0$ are Weibull shape and scale parameters, respectively. Furthermore, $\beta\geq1$ if the inference for the *i*^th^ barcode to represent a singlet improves as $b_{i}$ rises.

The closely related Fréchet probability function bounded at $x>0$ is suitable to represent the distribution of $b_{i}$ in the theoretical combinatorial index-based library, in which barcodes are assigned to single cells and accrue total UMI counts well over artifacts randomly spread throughout; it is defined as

$$\text{P}_{\text{C}}\left( b_{i}\leq x \right) \sim\text{Fr}\left( x;\mu,\nu\right)=\frac{\mu}{\nu}\cdot\left( \frac{x}{\nu} \right)^{-\mu-1}\cdot exp\left[ -\left( \frac{x}{\nu} \right)^{-\mu} \right]$$

in which $\text{P}_{\text{C}}$ refers to the probability function of a theoretical combinatorial index-based library, and $\mu>0$ and $\nu>0$ are Fréchet shape and scale parameters, respectively. Also, $\mu\geq1$ if the probability $\text{P}_{\text{C}}$ for any *i*^th^ barcode to approach the minimum value of $b_{i}>0$ is asymptotically small – i.e. $\text{P}_{\text{C}}$ speeds upwards at $b_{i}\gg0$.

Hence, assuming negligible error $\lim_{n\to\infty} \left( \varepsilon_{n}\text{δ}_{nk} \right)=0$, it can be shown the $\text{P}_{\text{C}}$-$\text{P}_{\text{D}}$ parametric mixture model for $b_{i}$ in a single-cell RNAseq library can be algebraically reduced to

$$\text{P}\left( b_{i}\leq x \right)=\left[ P_{D} \right]\cdot\left[ 1-w_{\text{C}}\cdot\left( 1+\text{P}_{\text{C}*\text{D}} \right) \right]$$

where

$$\text{P}_{\text{C}*\text{D}}=\frac{\Omega}{\text{A}}\cdot X^{\Omega-1}\cdot exp\left[ -\frac{1}{\alpha^{\beta}}\cdot\left( \frac{X^{\Omega}-\text{A}\cdot X}{\text{A}} \right) \right]$$

is a parametric probability distribution of the exponential family bounded at $\left\{ X=x^{\beta} \right\}>0$ and expressed in terms of composite parameters $\Omega=-\left( \mu/\beta\right)$ and $\text{A}=1/\left( \nu^{\mu}\cdot\alpha^{\beta} \right)$. Put simply, the $\text{P}_{\text{C}}$-$\text{P}_{\text{D}}$ mixture model predicts that, for any single-cell RNAseq library, the distribution of $b_{i}$ rises suddenly from a low-count detection limit defined by artifacts (consistent with a Fréchet distribution), exhibits a transition phase, tails off as $b_{i}$ values keep rising (consistent with a Weibull distribution), and stops as the expected values of $b_{i}$ reach an improbably high maximum value corresponding to multiplets.

The purpose of fitting a $\text{P}_{\text{C}}$-$\text{P}_{\text{D}}$ mixture model is to tease out which detected barcodes correspond to single cells based on their total UMI counts $b_{i}$. As mentioned earlier, we assume that single cells reside somewhere past the transition phase in the cumulative distribution of $b_{i}$ and into the tail-off range towards the maximum value of $b_{i}$. In that sense, one could infer – based on $b_{i}$ values alone – that barcodes representing single cells reside in an interphase $b_{\text{Low}}\leq b_{i}\leq b_{\text{High}}$ coalescing a Fréchet-dominated low-count domain ($b_{i}$ surges when going from artifacts to single cells) and a Weibull-dominated high-count domain ($b_{i}$ surges again when going from single to multiple cells). Thus, the expectation is that a $\text{P}_{\text{C}}$-$\text{P}_{\text{D}}$ mixture model will exist such that the underlying $\text{P}_{\text{C}}$ and $\text{P}_{\text{D}}$ distributions dominate at opposite extremes of the $b_{i}$ domain. If so, the scale parameters $\alpha$ and $\nu$ for $\text{P}_{\text{D}}$ and $\text{P}_{\text{C}}$, which denote the inflection points in each surge of the $b_{i}$ domain, can be used as fiducial marks to help infer limits in barcode coverages characteristic of single cells – i.e. $\nu\sim b_{\text{Low}}$ and $\alpha\sim b_{\text{High}}$. Accordingly, the shape parameters $\beta$ and $\mu$ estimate how steep those surges are when moving away from the $\nu$ and $\alpha$ fiducial marks, respectively.

To minimize the likelihood of admitting artifact barcodes when defining the single-cell interphase $b_{\text{Low}}\leq b_{i}\leq b_{\text{High}}$, we trim the $b_{i}$ domain inwards from the low-end fiducial mark $\nu$ to define $b_{\text{Low}}$. We reason that, being a Fréchet inflection point between noise and single-cell data, barcodes with $b_{i}=\nu$ are statistically indistinguishable from either. Thus, to prioritize certainty in single cell assignment, we impose a systematic shift such that $b_{\text{Low}}>\nu$. In doing so, we take three features into account: the relative Fréchet behavior contribution $w_{\text{C}}$, a combined steepness metric $\mu\cdot\beta$, and the proportional enrichment $\alpha/\nu$ between fiducial marks for the $\text{P}_{\text{C}}$-$\text{P}_{\text{D}}$ mixture model such that

$$b_{\text{Low}}=\nu\cdot\left( \frac{\alpha}{\nu} \right)^{\frac{w_{\text{C}}}{\mu\cdot\beta}}$$

corresponding, in logarithmic scale, to a weighed baseline shift from the low-end fiducial mark $\nu$ adjusted for steepness and not to encroach the high-end fiducial mark $\alpha$:

$$\ln\left( b_{\text{Low}} \right)=\ln\left( \nu\right)+w_{\text{C}}\cdot\left( \frac{1}{\mu\cdot\beta} \right)\left[ \ln\left( \alpha\right)-\ln\left( \nu\right) \right].$$

We recognize a similar case can be made at the high end of the $b_{i}$ spectrum – with some important modifications. In our interpretation of the $b_{i}$ probabilistic model, the transition going from artifacts to single cells reminisces a Fréchet-dominated probabilistic behavior. Thus, by fitting a single Frechét parametric distribution $\text{P}_{\text{Fr}}$ to the entire $b_{i}$ domain, while taking into account the contribution of low-count artefacts, such that

$$\text{P}_{\text{0}}\left( b_{i}\leq x \right) \sim\text{Fr}\left( x;\mu_{0},\nu_{0} \right)=\frac{\mu_{0}}{\nu_{0}}\cdot\left( \frac{x}{\nu_{0}} \right)^{-\mu_{0}-1}\cdot exp\left[ -\left( \frac{x}{\nu_{0}} \right)^{-\mu_{0}} \right]$$

one can project a parametrically determined $b_{\text{High}}$ single-cell upper bound for $\text{P}_{\text{0}}$ corresponding to a linear shift over the scale factor $\nu_{0}$ (a measure of the artefact-to-noise turning point), systematically weighed by the shape constant $\mu_{0}>1$, and mimicking a multiplet-free experiment:

$$b_{\text{High}}=\nu_{0}+2\cdot\mu_{0}\cdot\left( \overline{x_{\text{0}}}-\nu_{0} \right)$$

where $\overline{x_{\text{0}}}=\nu_{0}\cdot\Gamma\left( 1-1/{\mu_{0}} \right)$ is the expected arithmetic mean of $\text{P}_{\text{Fr}}$. Hence, our parametric definition of $b_{\text{High}}$ represents a doubling of the effective average gain in $b_{i}$ values among single cells vs. the baseline signal from artifacts, or $\left( \overline{x_{\text{Fr}}}-\nu_{0} \right)$, which we adjust by a factor of $\mu_{0}$ – i.e. the compactness of the histogram of observed $b_{i}$ values – to correct for projected dispersion in the $\text{P}_{\text{Fr}}$ distribution.

One important property of this definition for $b_{\text{High}}$ is that it is not only anchored – since fitting $\text{P}_{\text{Fr}}$ depends on the amount of artifact barcodes – but also self-ballasted: higher values of $\mu_{0}$ bring the tails of predicted $\text{P}_{\text{Fr}}$ histograms closer, shrinking the effective distance (or gain) between the $\nu_{0}$ baseline and the projected mean $\overline{x_{\text{Fr}}}$, while simultaneously stretching the linear shift between $b_{\text{High}}$ and $\nu_{0}$ by a factor of $\mu_{0}$. For example, even though values of the projected $b_{\text{High}}$ can vary widely in proportion to the baseline $\nu_{0}$ (between 3-6 times larger for typical $1.5<\mu_{0}<3$ fitted values) the actual rate of admitted barcodes, or $\text{P}_{\text{Fr}}\left( {b_{i}}/{\nu_{0}}\leq{b_{\text{High}}}/{\nu_{0}} \right)$, remains largely the same across the board (e.g. between 93^rd^-97^th^ percentile for typical $1.5<\mu_{0}<3$ fitted values). In other words, our parametric definition of the pass-fail criterion $b_{\text{High}}$ to distinguish singlets vs. multiplets is a frequentist projection that depends on two non-dimensional constants: $\mu_{0}$ and ${b_{\text{High}}}/{\nu_{0}}$. Combined, these properties predict two systematic advantages: a) the fraction of inferred singlets $b_{i}<b_{\text{High}}$ remains the same for a single-cell library even if re-sequenced to increase coverage, as artifact and singlet values both scale with $\nu_{0}$; and b) multiplets with $b_{i}\geq b_{\text{High}}$ are a systematic outlier subset, accounting for roughly the same fraction of detected barcodes overall, even for independent single-cell libraries that do not share the same fitted estimates for $\mu_{0}$ or $\nu_{0}$.

### References
