## Supplementary figures and images for "Patterns, Profiles, and Parsimony: dissecting transcriptional signatures from minimal single-cell RNA-seq output with SALSA"

### Figure S1

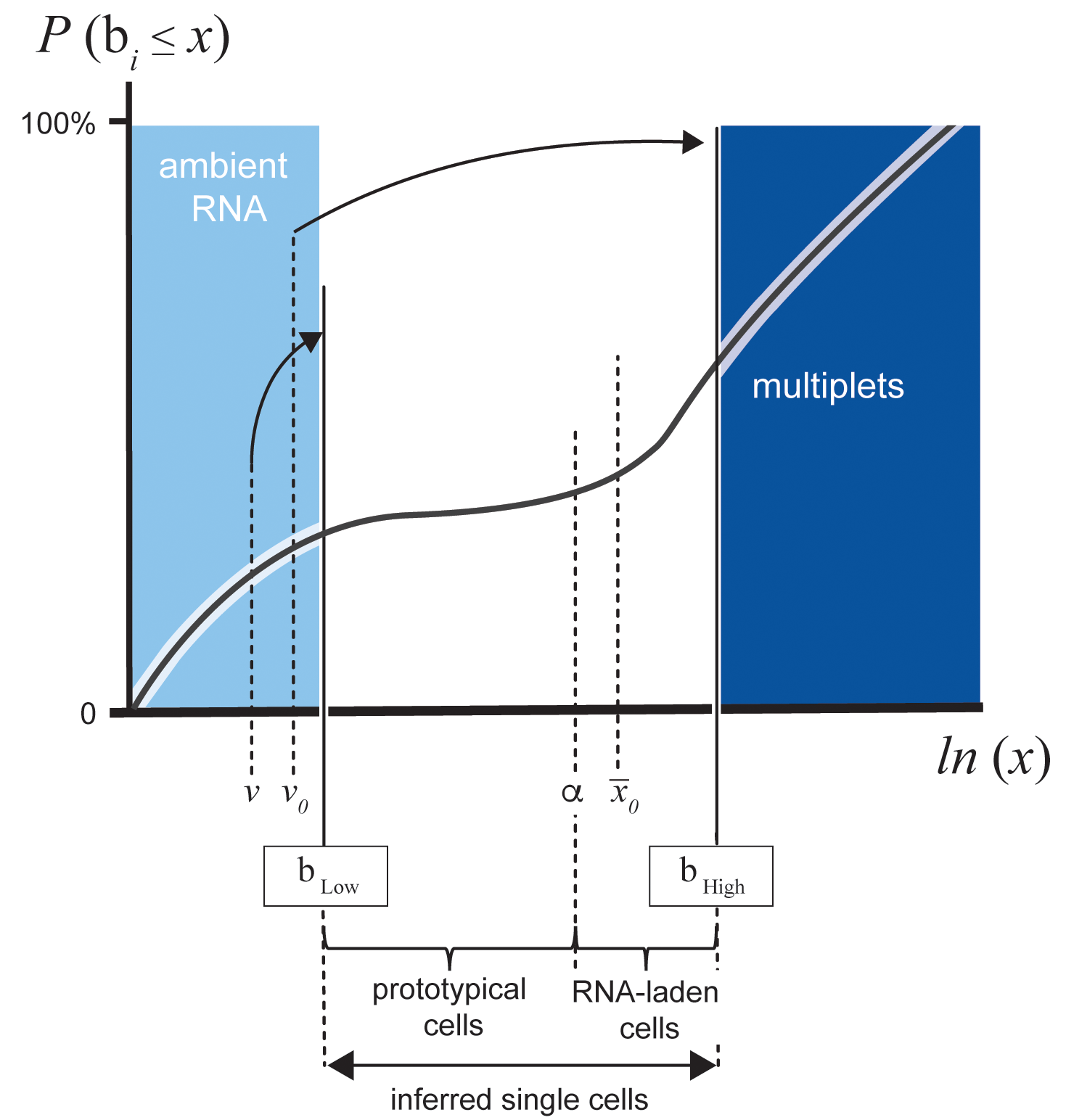

### Figure S2

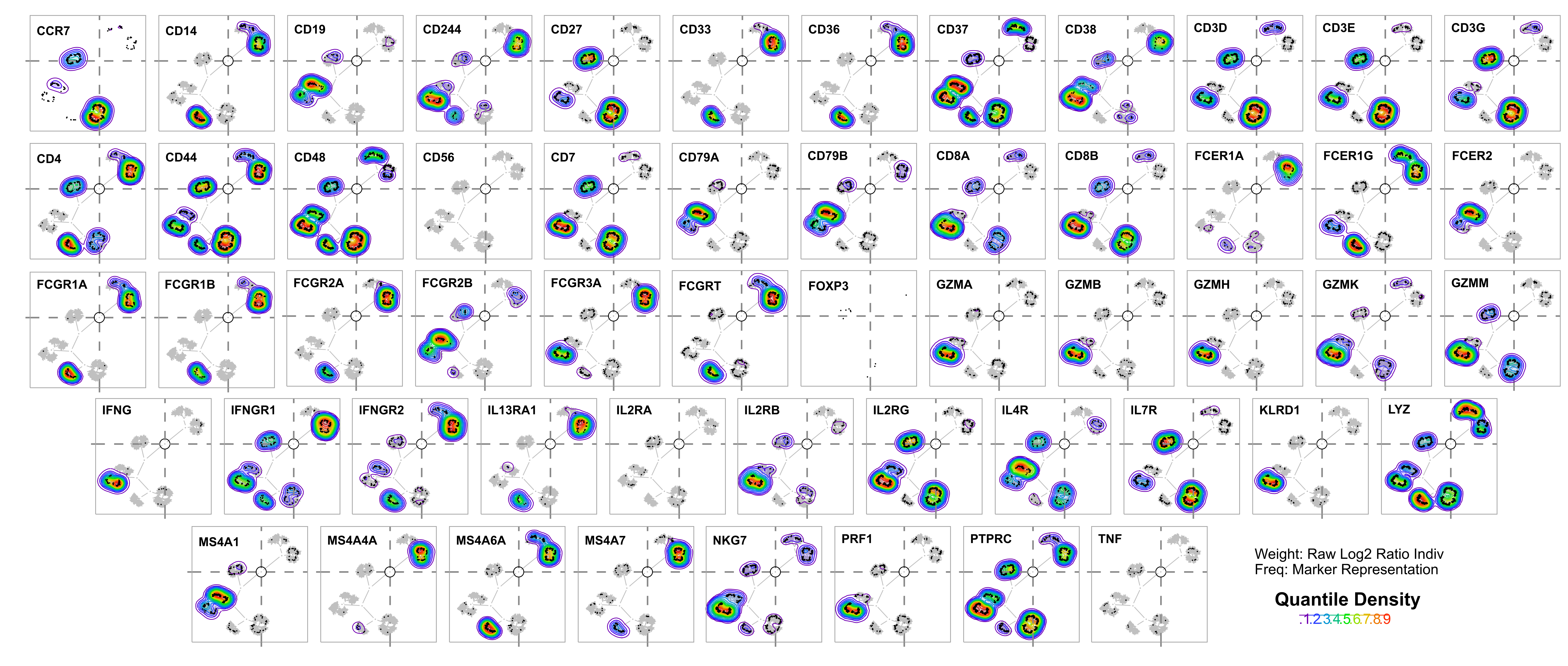

### Figure S3

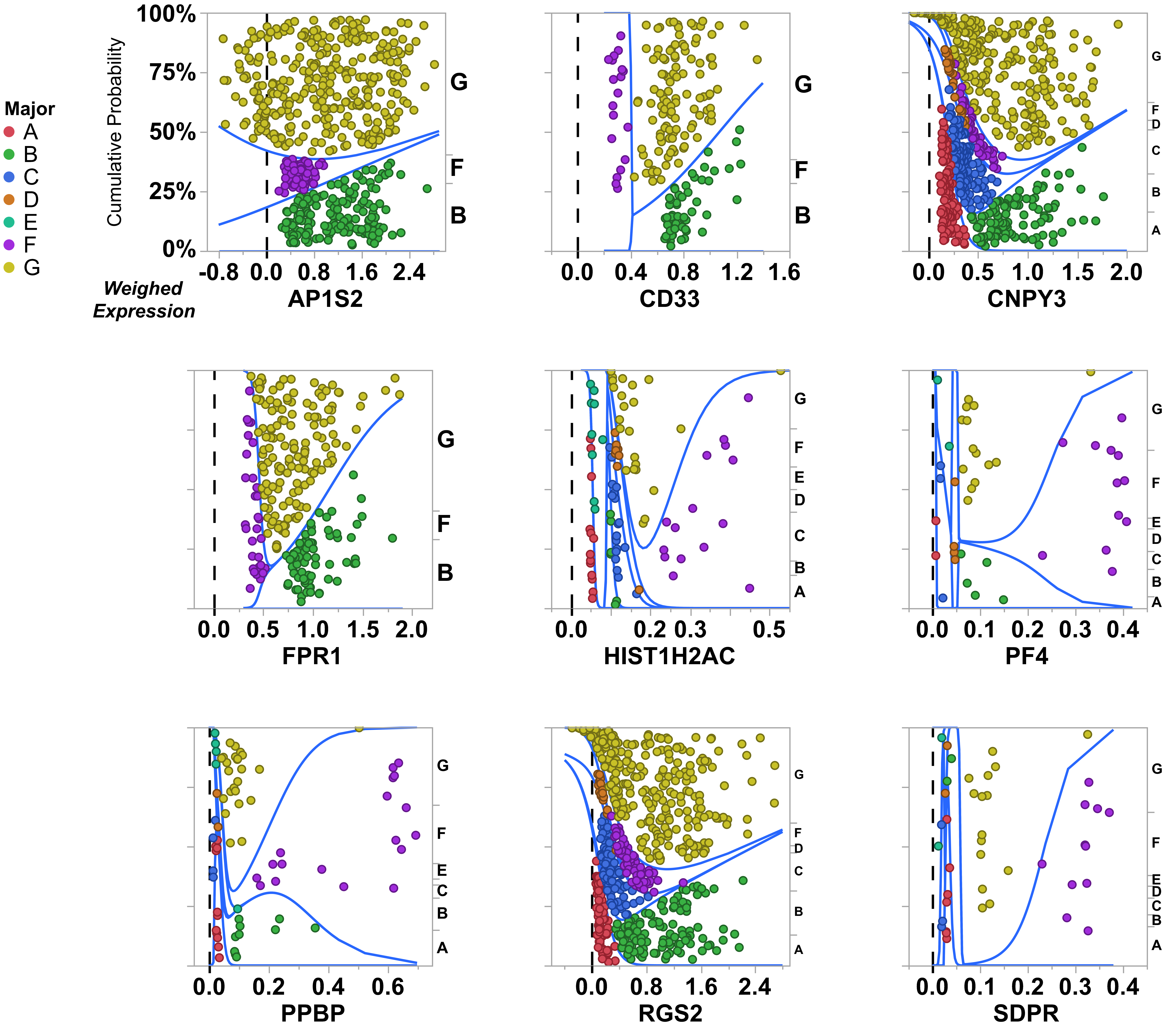
